# Dynamic linear models guide design and analysis of microbiota studies within artificial human guts

**DOI:** 10.1101/306597

**Authors:** Justin D Silverman, Heather Durand, Rachael J. Bloom, Sayan Mukherjee, Lawrence A David

## Abstract

Artificial gut models provide unique opportunities to study human-associated microbiota. Outstanding questions for these models’ fundamental biology include the timescales on which microbiota vary and the factors that drive such change. Answering these questions though requires overcoming analytical obstacles like estimating the effects of technical variation on observed microbiota dynamics, as well as the lack of appropriate benchmark datasets. To address these obstacles, we created a modeling framework based on **m**ultinomi**al l**ogistic-norm**a**l dynamic linea**r** mo**d**els (MALLARDs) and performed dense longitudinal sampling of replicate artificial human guts over the course of 1 month. The resulting analyses revealed that when observed on an hourly basis, 76% of community variation could be ascribed to technical noise from sample processing, which could also skew the observed covariation between taxa. Our analyses also supported hypotheses that human gut microbiota fluctuate on sub-daily timescales in the absence of a host and that microbiota can follow replicable trajectories in the presence of environmental driving forces. Finally, multiple aspects of our approach are generalizable and could ultimately be used to facilitate the design and analysis of longitudinal microbiota studies *in vivo*.

## INTRODUCTION

Artificial gut models have been used for decades to replicate the human intestinal environment and study the dynamics of resident microbes (1–3). These systems have the advantage of being sampled with arbitrary frequency, house environments that can be precisely controlled, and often face fewer ethical concerns than in-human studies (4). Artificial gut models have therefore been used to discover the effect of nutritional supplements on infant gut microbiota (5), mechanisms by how commensals repress *Salmonella* virulence (6), and dose-dependencies between microbiota-targeting therapies and metabolite production (7).

Yet, despite the utility and long development of artificial gut models, fresh questions regarding their fundamental biology remain. Such questions include the rapidity with which the composition of these microbial communities vary both in the presence and absence of perturbations (1, 8). While it is well known that *in vivo* microbiota may change on sub-daily time-scales due to host forcing, it is unclear whether such sub-daily dynamics may be seen in the absence of host effects (9). The degree to which replicate *ex vivo* systems exhibit stochastic behavior, or conversely behave deterministically, remains another outstanding question (1, 10). A deeper understanding of the reproducibility of artificial gut models has implications for our understanding of the relative importance of various factors in shaping the dynamics of host associated microbiota (11). For example, it is hypothesized that fixing the primary forces driving community composition would produce more reproducible microbiota dynamics than fixing secondary forces (12).

Gaining greater insight into the biology of artificial gut models though requires addressing analytical and statistical challenges. One key challenge is the often unquantified impact of artificial sources of intra-study variation such as variation due to sequencing counting and technical variation from sample processing (*e.g.*, unintended experimental errors and batch effects) (13–18). In particular, while the impact of variation due to sequence counting is more commonly addressed and modeled (13, 19–21), the impact of technical variation from sample processing is often unquantified and less well understood. Such technical variation may alter inference of both the magnitude and direction of variation among bacterial taxa. In addition, it is well established that sequencing studies of microbiota provide information only on the relative amounts of taxa and not their absolute abundance (22–24). The analysis of such relative (or compositional) data remains an open area of study and naive analyses can lead to a distorted view of the patterns of variation present in a community (21, 22, 25–27).

The design of *ex vivo* microbiota studies is also confronted by the lack of appropriate benchmark datasets. For example, defining suitable sampling frequencies for artificial gut studies requires insight into the timescales on which human gut microbiota fluctuate (28). Yet, even though gut microbiota dynamics *in vivo* and bacterial mock communities *in vitro* are known to behave on the timescale of hours (9, 29, 30), most longitudinal studies to date in *ex vivo* models are only sampled on the order of days (1, 8, 10, 31). Additionally, while the collection of technical replicates could be used to quantify the effects of technical variation (32), such replicate sampling is generally not performed in longitudinal microbiota studies.

Here, we integrated model development and experimental design to address key challenges facing the analysis and design of longitudinal artificial gut studies. We collected longitudinal samples with up to hourly frequency from replicate artificial gut models over the course of one month. We combined this longitudinal sampling with the collection of technical replicates so that we could characterize the impact of technical sources of variation on observed microbiota dynamics. To isolate separate biological and technical sources of community variation in our dataset, we created a modeling framework called MALLARD that is based on a class of generalized dynamic linear models appropriate for microbiota time-series data. Together, our dataset and modeling framework allowed us to investigate the patterns and time-scales of microbiota variation in an artificial human gut.

## RESULTS

### Longitudinal modeling

To separate biological and technical variation in artificial gut time-series, we introduce an extension of Dynamic Linear Models (DLMs) tailored for microbiota data. DLMs have widespread use including industrial applications such as commercial forecasting and engineering control systems (33). At their core, DLMs model a system as a time-varying state that is observed through a noisy process. We extended DLMs to a class of **<u>m</u>**ultinomi**<u>al</u> <u>l</u>**ogistic-no**<u>r</u>**mal **<u>d</u>**ynamic linear models by building off of the work by Cargnoni, Muller and West (34). We refer to this as the MALLARD class of models.

We analyzed the artificial gut dataset described below using a MALLARD model that is generative and assumes there exists an unobserved microbial composition (*θ_t_*; the state) that evolves through time (Fig. 1A) due to stochastic biological variations (*w_t_*). We regard the state sequence (*θ_1_, …,θ_t_,…,θ_T_*) as the true microbial dynamics in a time-series. Random technical variations (*v_t_*) are then added to the true system state (*θ_t_*)resulting in the composition *η_t_* (Fig. 1B). We observe *η_t_* through a multinomial counting process. This formulation is similar to the constant level model commonly used in Bayesian time-series analysis (34). By separately modeling the process generating *w_t_* and *v_t_* with distinct covariance matrices (*W* and *V*, respectively), we can decouple biological and technical variations in artificial gut datasets (Fig. 1C). Visually, we found this model provided a good fit to artificial gut data (Fig. S1).

**Figure 1.**
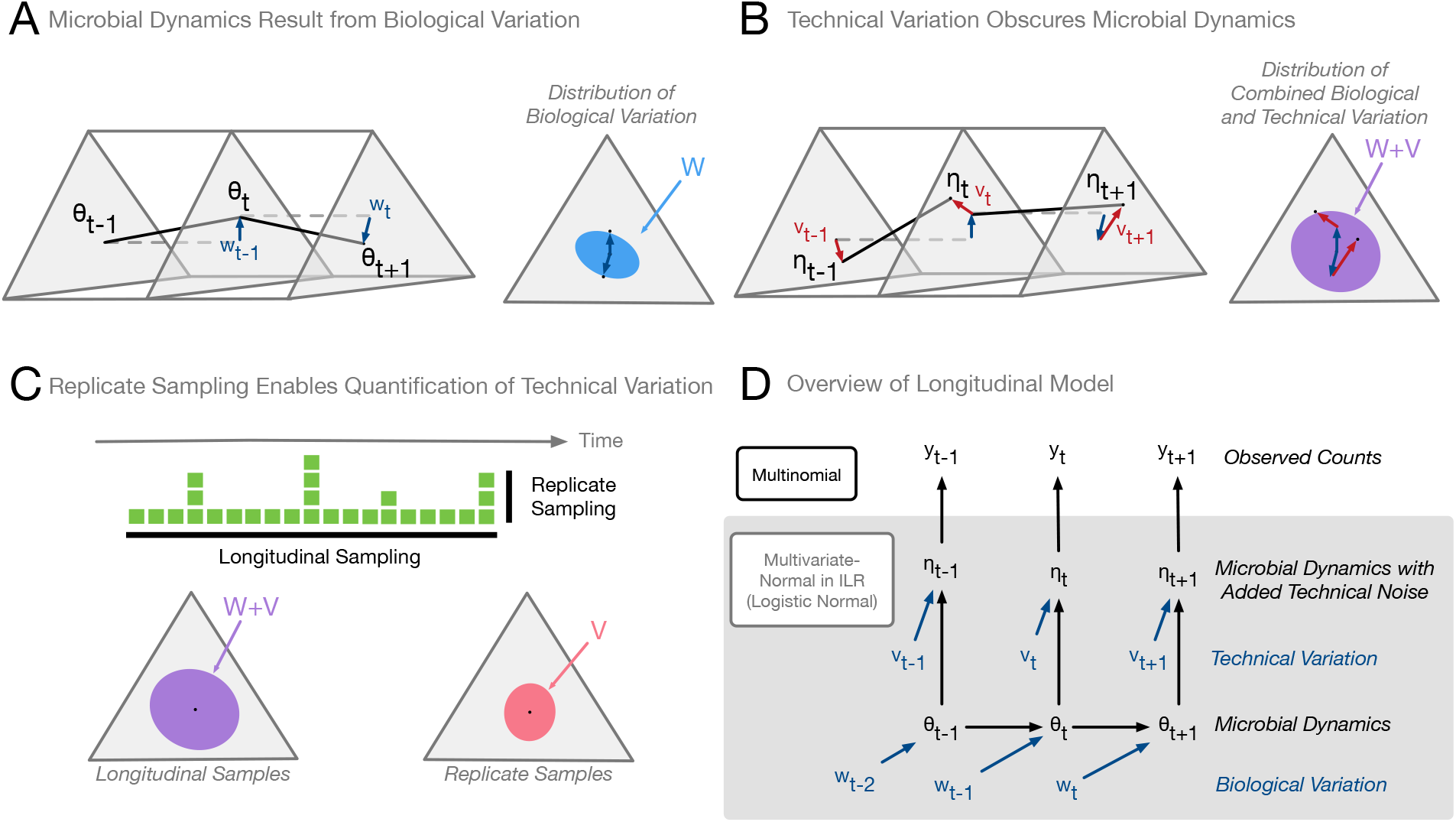
A generative model for microbial dynamics obscured by technical variation. (A) Microbial dynamics result from biological variation. The time series (*θ_1_,…,θ_t_, …,θ_T_*) defines the dynamics of a microbial community and results from biological variations (*w_1_, …, w_t_, …, w_T_*) which are assumed to be independent and identically distributed (i.i.d.) logistic-normal with mean zero and covariance *W*. (**B**) **Technical variation obscures microbial dynamics**. Technical variation (*v_1_, …, v_t_, …, v_T_*) from sample processing introduces noise into measurements of microbial dynamics and are assumed to be i.i.d. logistic-normal with mean zero and covariance *V*. (**C**) **Replicate sampling enables quantification of technical variation**. Hypothetical collected samples are denoted by green squares. Differences between longitudinal samples reflects both biological and technical sources of variation. In contrast, differences between technical replicates (samples from the same time point) should reflect only technical sources of variation and can be used to estimate *V* (*Methods*). (**D**) **Overview of longitudinal model**. The microbial dynamics and the confounding technical variation (A and B) are modeled in a ILR space such that the logistic-normal distribution transforms to a multivariate-normal distribution for mathematical convenience (*Methods*). The observed count data is assumed to be distributed multinomial from the compositions (*η_1_, …, η_t_,…,η_T_*) (*Methods*).

### Daily and hourly gut microbiota time-series in an artificial human gut

We applied our model to an artificial gut that was constructed using continuous-flow anaerobic bioreactor systems that have been validated as models of human gut microbiota (1, 6, 15, 35, 36). The same starting human fecal inoculum was seeded into replicate *ex vivo* vessels (*n*=4) and cultured for 1 month (Fig. 2 and S2). Throughout the experiment pH, temperature, media input rates, and oxygen concentration were all fixed (*Methods*). To introduce microbial dynamics into our systems, a single bolus of *Bacteroides ovatus* isolated from the stool donor was administered to the system on Day 23 (*Methods*). The *B. ovatus* bolus did not have discernable effects on microbial dynamics, but the media it was suspended in appeared to induce minor shifts in the relative abundances of select bacterial taxa that were visible with hourly sampling (Fig. S3). Additional microbial dynamics related to media input were generated by an inadvertent feed disruption in two vessels between Days 11 and 13 of the study. Overall, similar to previous studies (1, 10), the artificial gut maintained much of the microbial diversity present in the inoculating stool: 91% of bacterial families present Days 1-5 of the study were detected between Days 23-28. The 9% of bacterial families that did not persist represented only 0.06% of the total sequencing reads in the dataset.

**Figure 2.**
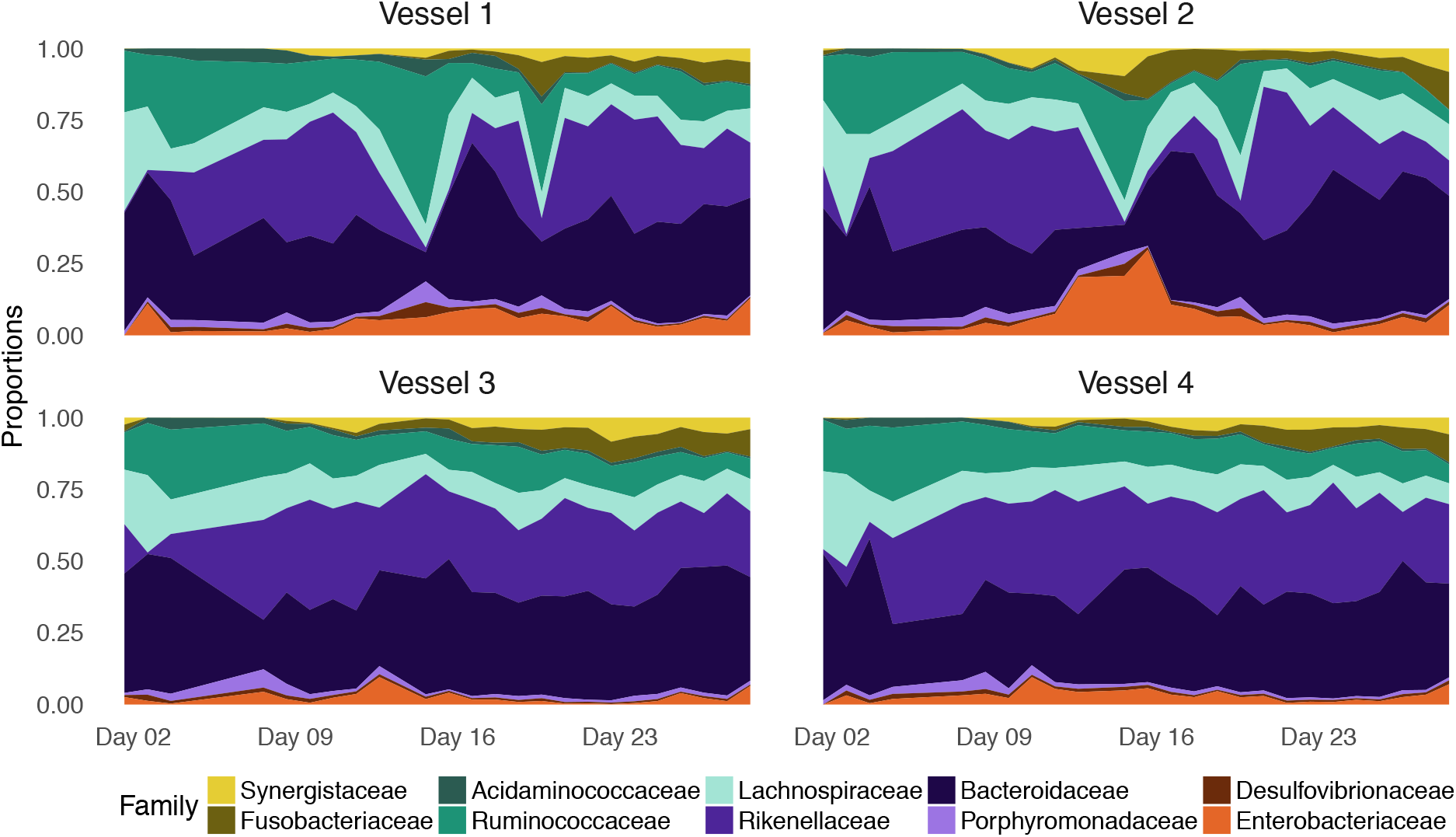
Proportions of 10 most abundant bacterial families over time. Four identical continuous-flow anaerobic bioreactor systems were each inoculated from a single human fecal specimen and cultured over the course of 1 month. Proportions of bacterial taxa at sampled time points were estimated by dividing observed read counts from 16S rRNA sequencing by the total number of counts observed for each sample. In addition to daily sampling (shown here), hourly samples were taken from each vessel (480 hourly samples total) as well as 20 technical replicate samples from the final time point of each vessel (Fig. S4). Proportions from the hourly sampling data are shown in Figure S3.

To investigate the timescales on which microbial communities vary *ex vivo* and to characterize the impact of technical variation on our measurements, we developed a sampling scheme for our artificial gut that featured fine temporal resolution and numerous technical replicates (Fig. S4). As in previous studies, all four vessels of the artificial gut were sampled daily over the course of one month (Fig. S4). To investigate potential sub-daily variation, we aimed to oversample the system and collected 120 sequential hourly samples, from each replicate vessel, during a five-day period. To estimate the technical variation in our measurements, we collected 20 replicate samples from the final time point of each artificial gut vessel (Fig. 1C and S4; *Methods*).As samples collected in the same vessel at the same time are expected to be biologically identical (*i.e.*, feature no biological variation), any variation in the resulting community measurements between these technical replicates can be ascribed solely to technical sources (Fig. 1C). To ensure that the technical variation profile of the replicate sampled matched that of the longitudinal samples, all samples were randomized and processed together for sequencing.

### The structure and magnitude of technical variation from sample processing

After fitting the MALLARD model to the resulting artificial gut dataset, we investigated the technical variation from sample processing (*V*) to our inference of variation due to biological sources (*W*). Variation from sample processing and variation due to sequence counting were represented as separate processes (Fig. 1D). Differing correlation structure between *V* and *W* would support our hypothesis that sources of technical variation could obscure biological forces acting on microbiota within an artificial gut. Indeed, a permutation analysis indicated it highly improbable that *V* and *W* had the same correlation structure (posterior probability < 1%; *Methods).* Lowdimensional projections of the posterior distributions of *V* and *W* supported this conclusion and revealed how overall variation patterns involving bacterial families like Lachnospiraceae, Fusobacteriaceae and Bacteroidaceae more strongly resembled patterns of technical variation than biological variation (Fig. 3A). Thus, some patterns of covariation among taxa in artificial gut studies may be due to technical sources of variation.

**Figure 3.**
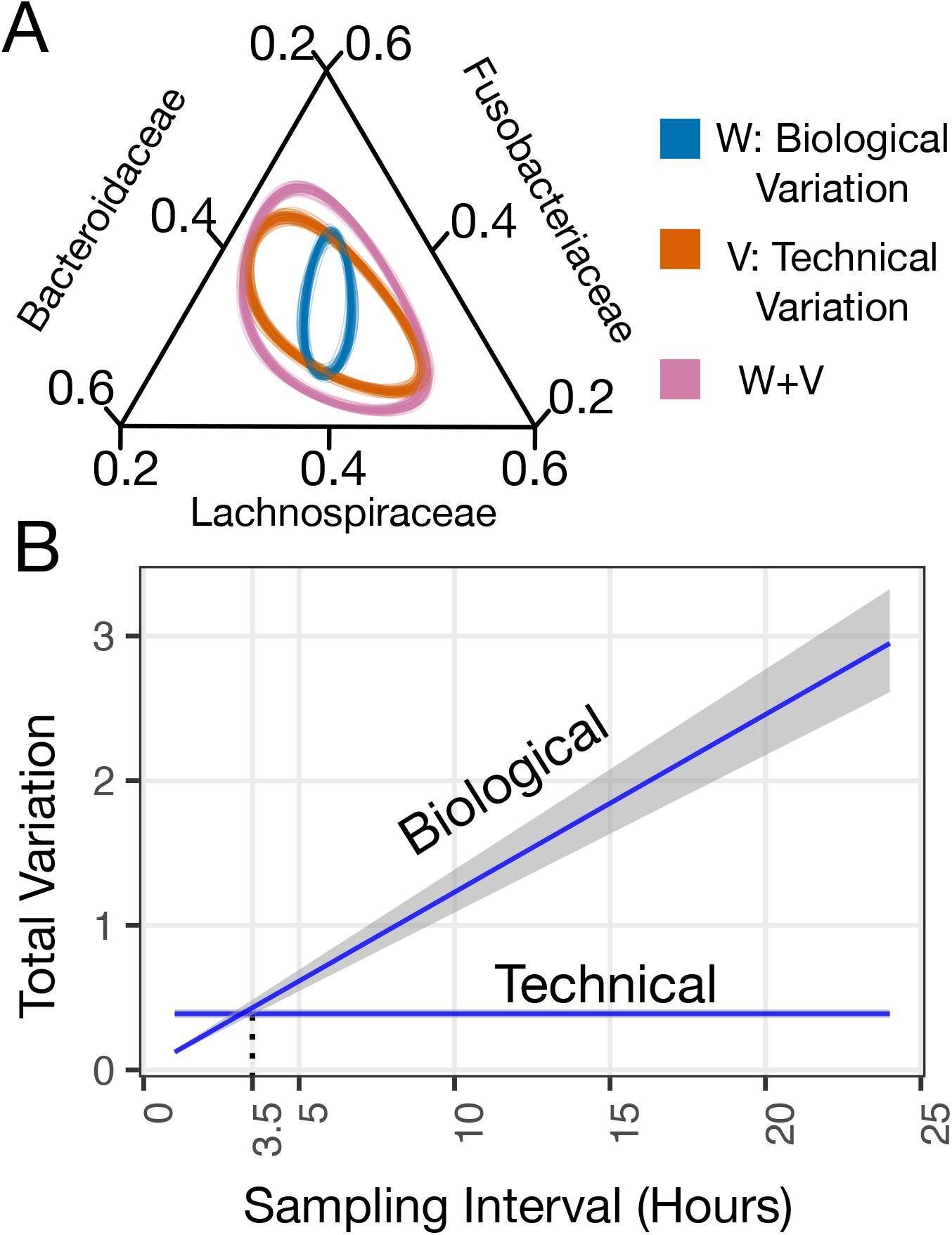
Structure and magnitude of biological and technical variation. (**A**) A two-dimensional projection of 100 samples from the posterior distributions of *W* and *V* inferred from the artificial intestine time-series displayed as a ternary diagram (centered 95% credible regions of the logistic-normal distribution shown). (**B**) Mean and 95% credible interval for total biological (*Tr*(*W*)) and total technical (*Tr*(*V*)) variation as a function of sampling interval in hours.

We next investigated the relative magnitude of technical variation from sample processing and biological variation as a function of sampling frequency. Total variation is defined as the trace of a covariance matrix (37). We therefore computed *Tr(W)/Tr(V)*, which is analogous to the signal-to-noise ratio used in signal processing (38). We found that for our time-series analyzed on an hourly basis (*Methods*), the median ratio of biological to technical variation was 0.30 (0.26-0.36 95% credible interval). We therefore estimate that only 24% (21%-26%) of the total variation in relative abundances was due to biological sources. This result was insensitive to perturbation of our priors (Fig. S5). Moreover, because variance is additive between time points within our model, we could estimate biological and technical variation as a function of sampling interval (Fig. 3B; *Methods*). We found that at a sampling interval of 3.5 hours, the total biological and technical variation were approximately equal. At sampling frequencies faster than 3.5 hours, technical variation outweighed biological variation; at slower sampling frequencies, signal associated with biological variation exceeded technical noise.

### Timescales of microbial dynamics

In order to explore the time-scale of microbial dynamics within replicate artificial gut vessels, we visualized microbial dynamics using PhlLR balances. PhlLR balances represent the log-ratio between phylogenetically neighboring clades of taxa, providing a phylogenetically informed way to study microbial dynamics without compositional artifacts (22). We quantified the magnitude of changes in balance values in units of evidence information (*e.i.*), which are a measurement of compositional change (39). A 1 *e.i.* change is equivalent to an approximately 4-fold change in the ratio of two bacterial taxa and a 2 *e.i.* change is equivalent to an approximately 17-fold change in the ratio of two taxa (see *Methods* for further discussion). Multiple balances exhibited sub-daily dynamics (Fig. S6,7), most notably the ratio of bacteria from the phylum Bacteroidetes to bacteria from the phyla Proteobacteria and Fusobacteria (Fig. 4). The balance between Bacteroidetes and Proteobacteria/Fusobacteria appeared to fluctuate on timescales shorter than 1 day with an amplitude of approximately 1 *e.i.* (0.5-1.5, 95% credible interval; Figure 4B). Balance dynamics did not correspond to recorded environmental or technical variations (*e.g.* media changes, identity of the researcher collecting the sample, sequencing batch number, *B. ovatus* supplementation, or the feed disruption of vessels 1 and 2) and did not display an exact 24-hour periodicity (Fig. S8). Balance fluctuations were observed in all four replicate artificial gut models, but did not appear to be synchronized as would be expected if these dynamics were driven by a shared environmental factor (Fig. S7). We ultimately could not identify a technical or environmental cause of fluctuating balance dynamics on sub-daily timescales.

**Figure 4.**
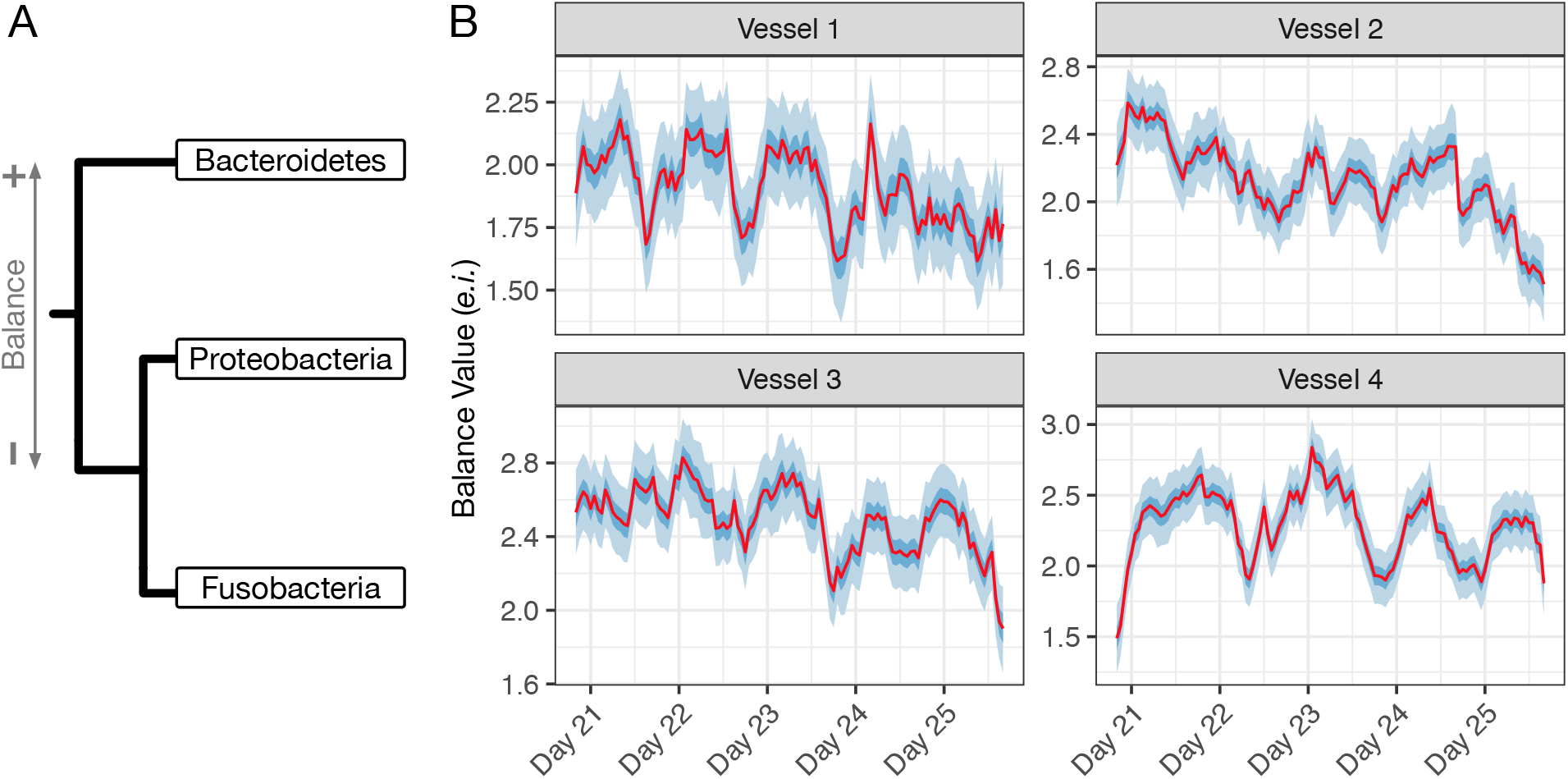
Sub-daily microbiota dynamics within the artificial intestine. (**A**) An annotated phylogenetic tree defining PhILR balance between the phyla Fusobacteria, Proteobacteria, and Bacteroidetes. The balance reflects the scaled log-ratio of the geometric mean relative abundance of families in the Bacteroidetes phyla (numerator, +) to the geometric mean relative abundance of families in the Fusobacteria and Proteobacteria phyla (denominator, −) (*Methods*). (**B**) Mean, 50%, and 95% credible interval of posterior distribution for *θ* (microbial dynamics in absence of technical variation) for the PhILR balance defined in (A). The full list of PhILR balances and corresponding posterior estimates are shown in Figure S6 and S7.

### Exploratory Analysis of Biological Variation

We next explored the biological variation captured in our model by investigating the pairwise variation between bacterial families in our dataset. Direct taxon-level analysis of the covariance matrix *W* is difficult though because the elements of this matrix are balances - not individual taxa. We therefore investigated the temporal variation of specific taxon pairs using a tool from Compositional Data Analysis called the variation array (40). A variation array represents the variance of the log-ratio between pairs of taxa (*Methods*). When variance of a pairwise log-ratio is near zero the two taxa positively covary; whereas when this variance is high the two taxa exhibit either unlinked or exclusionary patterns (22, 41). The resulting variation array (corresponding to *W*) revealed that most temporal variation was contained in log-ratios between four bacterial families: the Rikenellaceae, Synergistaceae, Enterobacteriaceae, and Fusobacteriaceae. By contrast, log-ratios between other bacterial families were approximately 1 order of magnitude smaller (Fig. 5A). Overall, we estimated that the Rikenellaceae, Synergistaceae, Enterobacteriaceae, and Fusobacteriaceae accounted for 72% (69% - 75%; CLR basis) of the total biological variation seen in the dataset (Fig. S9).

**Figure 5.**
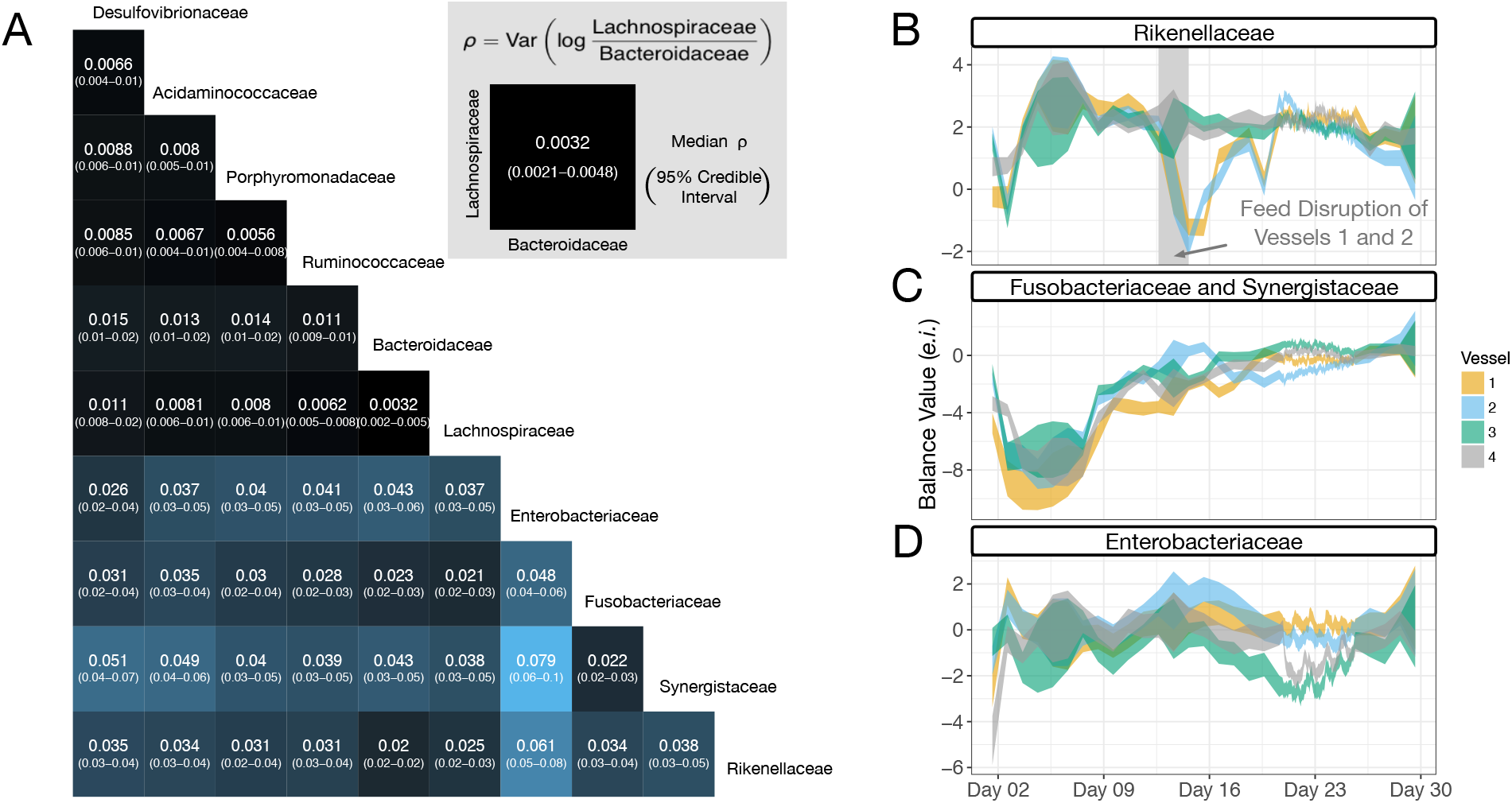
The decomposition of biological variation among bacterial families. (A) Heatmap of posterior distribution of log-ratio variance (*ρ*) between pairs of bacterial families. Heatmap color is given by the median of the posterior distribution of *ρ*. Columns and rows refer to the bacteria in the numerator and denominator of the corresponding log-ratios respectively. A similar decomposition of technical variation is shown in Figure S10. (**B-D**) The highest variance bacteria displayed distinct temporal patterns (95% posterior credible regions for *θ*). (**B**) The balance between the family Rikenellaceae versus all other bacterial families decreased substantially during the feed disruption of vessels 1 and 2. (**C**) The balance between the Fusobacteriaceae and Synergistaceae versus all other bacterial families slowly increased over the course of the study after an initial acclimation period. (**D**) The balance between the family Enterobacteriaceae versus all other bacterial families displayed fluctuating dynamics likely related to those shown in Figure 2D.

We noticed an inverse relationship between the biological variation of taxa to their relative abundance in the starting inoculum. We observed this relationship by fitting a linear regression model to each posterior sample in a CLR-transformed space (95% posterior credible interval for regression slope: −0.26 to −0.18; Fig. S11; *Methods*). Thus, our analyses suggested that more variable taxa in the artificial gut tended to be rarer at the time of inoculation. This result was insensitive to perturbation of our prior assumptions (Fig. S12), and there was only a 3.8% posterior probability that the observed inverse relationship was an artifact of low abundance families being more numerous than high abundance ones (*Methods*).

To better understand the patterns of variation we observed from the above analysis, we manually created three balances which together highlighted patterns that we believe explain the high variability of the Rikenellaceae, Synergistaceae, Enterobacteriaceae, and Fusobacteriaceae families (Fig. 5B-D). Together these three manually curated balances explained 73% of the biological variation in the dataset (69%-76%, 95% credible interval). This manual curation was informed by the inspection of microbial dynamics within the PhILR basis and a hierarchical cluster analysis which suggested that the Fusobacteriacae and Synergistaceae shared similar dynamic patterns and should therefore be grouped together for analysis (*Methods*, Fig. S13). One highly variable balance, which represented the ratio of Rikenellaceae to the other 9 bacterial families, exhibited a 3 *e.i.* decrease during Days 11-13 in two artificial gut replicate vessels before recovering over the subsequent week (Fig. 5B). This decrease corresponded with the transient feed disruption in those replicate vessels. A second highly variable balance, the ratio between the Fusobacteriaceae and Synergistaceae to all other bacterial families, increased by approximately 8 *e.i.* between Days 2-7 and the end of the experiment (Fig. 5C). The nadir of this balance coincided with an initial microbiota adaptation phase that is repeatedly reported in artificial gut studies (1, 10). A third balance, representing the ratio of Enterobacteriaceae to the other 9 bacterial families, exhibited irregular sub-daily oscillatory behavior (Fig. 5D and S14). These balance dynamics are likely related to those observed earlier between the phyla Bacteriodetes and Fusobacteria/Proteobacteria (Fig. 3), given the membership of Enterobacteriaceae within the phylum Proteobacteria.

## DISCUSSION

Here, we created a modeling framework that allowed us to partition a densely sampled artificial human gut time-series into components associated with technical and biological sources of variation. Our results demonstrate that technical source of variation influence *ex vivo* microbiota dynamics, accounting for 76% of community variation on hourly time-scales. Still, we observed evidence for bona fide microbiota dynamics on sub-daily timescales. Our integrated analyses also resulted in approaches for characterizing microbiota variation over time. By investigating the distribution of biological variation among taxa we identified three distinct dynamic patterns that accounted for over 70% of the variation within our dataset. Together, our results contribute to our understanding of the dynamics of *ex vivo* artificial gut systems, as well as the design and analysis of future longitudinal microbiota studies.

Our findings in concert offer several insights that could be useful for the design and analysis of *ex vivo* studies of human gut microbiota. Our study suggests that technical variation introduced during sample processing can affect observations of microbiota dynamics not just in scale but also in the patterns of covariation among taxa. To date, knowledge has been limited on the effects of technical variation in *ex vivo* studies of human gut microbiota. Select exceptions include prior bioreactor studies that found technical variation to be limited and dominated by biological variation (10).

Our methods and results also have implications for the choice of sampling frequency in *ex vivo* artificial gut studies. While oversampling can waste resources, under-sampling can bias inference through a mechanism known as signal aliasing (28, 42, 43). Here, to balance between resource-intensive exploration of sub-daily dynamics while still capturing longer multi-day dynamics, we made use of MALLARD and mixed rate sampling (*e.g.*, sampling at both daily and hourly timescales). Our findings suggest that future studies interested in tracking the full range of *ex vivo* microbiota dynamics or that focus on community responses to rapidly changing external factors (e.g. single dose small molecule or prebiotic supplementation studies), should consider collecting sub-daily measurements. By contrast, studies interested solely in longer term changes (e.g. overall nutritional studies), may find daily sampling sufficient. Additionally, by combining replicate sampling with purposeful oversampling, we were able to determine an effective signal-to-noise ratio as a function of sampling frequency. This ratio could be used to estimate an upper-limit sampling frequency above which the benefits of increases samples are diminished by the relatively high levels of technical variation.

Another methodological approach we develop here for artificial gut studies is an exploratory technique for discovering temporal patterns among microbial taxa. As gut microbiota time-series often involve many taxa, methods of dimension reduction can aid data interpretation. Yet, many standard time-series tools for dimension reduction, such as dynamic principle component analysis, are not well suited for microbiota data in the form of relative taxa abundances (37). We overcome this challenge and identify distinct patterns in our dataset using a tool from compositional data analysis called the variation array (40). Hierarchical cluster analysis and manual curation then allowed us to identify three balances (Fig. 5B-D), highlighting four bacterial families, that together accounted for over 70% of the variation in our dataset. These three balances revealed key dynamical patterns and aided in our interpretation of the forces acting on our artificial intestine.

Beyond methodology for designing *ex vivo* gut microbiota studies, our findings also have several implications for the underlying biology of these systems. First, we observed distinct differences in the replicability of dynamics between vessels at hourly compared to daily time-scales. In keeping with previous reports, we found that replicate artificial guts display replicable dynamic patterns when analyzed on daily timescales (1). Notably, even after a feed interruption, vessels 1 and 2 returned to a similar composition as the two continuously fed vessels after roughly one week and appeared to display similar overall dynamics thereafter (Fig. 5B and S6). Such observations suggest that these systems follow distinct and potentially predictable trajectories on the time-scales of days. Conversely, dynamics appeared less synchronized at hourly time-scales (Fig. S7). For example, balances involving the family Enterobacteriaceae (*e.g.*, balance n12), which demonstrate oscillatory patterns in multiple replicate vessels, do not appear in phase with one another. Together, these observations suggest a conceptual model for our artificial gut systems in which strong deterministic forces such as resource availability drive predictable community dynamics on longer time-scales while other, potentially more subtle and variable ecological forces, drive temporal variation on shorter time-scales.

Another implication of our results to the biology of *ex vivo* gut microbiota involves sub-daily dynamics of bacterial families such as the Enterobacteriaceae. We acknowledge that despite our efforts to keep culture conditions constant in the artificial gut, we cannot exclude the possibility that these dynamics were caused by fluctuating and unmeasured aspects of our artificial gut setup (*e.g.*, room temperature, ambient humidity, room lighting). Although, we consider it unlikely that such external driving forces would cause dynamics that were unsynchronized between replicate vessels (Figure S7). Moreover, rhythmic dynamics within microbial communities have previously been observed or predicted in host-free environments (44–46), and bacteria have been speculated to harbor circadian clocks (47). The dynamics we observed may have even underestimated the rate at which microbiota varied in our system as genomic material likely remains measurable for a period after cell death and our analysis at the family level could smooth dynamics at finer taxonomic scales. Thus, our results support hypotheses that dynamical properties inherent to microbiota could contribute to subdaily oscillations observed in the mammalian gut (48).

We also found that rare bacterial taxa tended to harbor the greatest variation over time. Such variation may reflect the expansion of dormant enteric microbes that take advantage of differences between *ex vivo* and *in vivo* environments to expand (49, 50). In particular, we found that in all four artificial gut vessels the Fusobacteriaceae and the Synergistaceae underwent an approximately 4 e.i. drop in their relative abundance upon transplantation from the *in vivo* to *ex vivo* environment and then a subsequent increase to approximately 4 *e.i.* greater than their initial levels (Fig. 5C). Such patterns may reflect a form of ecological succession in which these bacterial families are well suited for the *ex vivo* environment but are initially outcompeted by faster growing microbes. Such succession is well described in both *in vivo* and environmental ecosystems (51–53), and supports the general hypothesis that select theoretical frameworks from the field of environmental microbial ecology are also applicable to host-associated microbiota (54).

Still, insights from our study here into the methodology and underlying biology of artificial gut experiments have several limitations. First, some sources of technical variation, such as systematic biases introduced due to separation of samples into batches for DNA extraction, PCR, or sequencing may have exploitable and reproducible structure that could be controlled for during modeling thus improving the resolution of longitudinal microbiome studies. Here, we have chosen to treat these sources of variation as random *en batch* and instead leave such extensions of our analysis for future work. We note however, that mixed effects models, which have proven to be a powerful method of accounting for such systematic biases or batch effects, and their dynamic extensions both represent special cases of dynamic linear models (38). We therefore believe MALLARD will prove useful in modeling and controlling for such batch effects. Second, computational limitations restricted our analysis to relatively few dimensions. Due to this limitation, we analyzed only the 10 most abundant bacterial families, which in turn could have masked dynamics occurring at finer taxonomic scales (28). Two future refinements may improve computational efficiency: univariate filtering methods for multivariate time-series have been developed and would reduce the dimensionality of matrix manipulations used in MALLARD (55); and, emulation methods that iteratively refine probabilistic models have been developed for high-dimensional count data and show promise (56). Third, we chose to collect all technical replicates from the same time-point and relied on randomizing both longitudinal and replicate samples into batches to ensure that the replicate samples faithfully represented the technical variation profile in our study. As an alternative, we could have instead spread technical replicate samples out over the course of the experiment, collecting duplicate or triplicate samples at regular intervals. As the MALLARD framework enables technical replicate samples to be collected with any distribution within the study, distributed replicate samples may have estimated microbial dynamics with differing precision.

Despite our study’s limitations though, we believe our methodological insights useful for the design of artificial gut experiments; and, more broadly, designing *in vivo* longitudinal studies of host-associated microbial communities. Longitudinal studies in humans have provided unique insights into disease and therapy (28, 43), including optimal antibiotic treatment regimens (57), mechanisms underlying disease and recovery from acute secretory diarrhea (58) or intestinal cleanout (59), as well as identifying vaginal microbiota signatures associated with preterm birth (60). Aspects of our modeling framework here could be applied to future longitudinal analyses in humans. Specifically, attention has been given recently to issues arising in the analysis of relative microbiome data (22–27, 41, 61, 62) and there is growing awareness of how count variability influences microbiota surveys (20, 21, 63). Missing data are also a common challenge in longitudinal microbiome analyses, particularly when the temporal evolution of observed data is modeled. MALLARD is unique in working at the intersection of compositional analysis, count variability, and linear longitudinal models that can account for technical variation in datasets with many missing observations (64–67).

Aspects of our artificial gut sampling design could also be applied to longitudinal microbiota studies in humans. Replicate sampling could be used to quantify the effects of technical variation for *in vivo* microbiota time-series. Moreover, choosing an appropriate sampling frequency remains an outstanding challenge in translational microbiome studies (68). Pilot time-series experiments that intentionally sample *in vivo* systems more frequently than expected microbiota dynamics (*i.e.*oversampling) could be used to compute tradeoffs between technical and biological variation at higher sampling frequencies as we do here; aiming to sample at frequencies where technical variation does not exceed biological variation could help guide economical use of laboratory resources. Indeed, potentially oversampled longitudinal datasets for human gut microbiota already exist (albeit with limited replicate technical sampling) (69, 70). Ultimately, by improving the design and analysis of microbial community dynamics in longitudinal *in vivo studies*, an improved understanding of the role of microbiota in human health and disease can be achieved.

## METHODS

### ARTIFICIAL INTESTINE EXPERIMENTS

#### Collection and Preparation of Fecal Inoculate

A fresh fecal sample was obtained from a healthy volunteer who provided written informed consent (Duke Health IRB Pro00049498). The sample was stored and prepared for inoculation into an artificial gut systems within an anaerobic chamber (Coy). The fecal sample was weighed into 50ml conical tubes, approximately 5 grams per tube, and then pre-reduced MGM (McDonald Gut Media; (1)) was used to fill the tube. The fecal matter was homogenized briefly using a benchtop vortex and then centrifuged 10 min at a speed of 175 × g. The supernatant was decanted into syringes for inoculation.

#### Artificial Gut Preparation

A four-vessel continuous flow artificial gut system (Multifors 2, Infors) was used to culture gut microbiota seeded from human stool. Vessels were sterilized and prepared with 300 ml of fresh MGM. Inoculation of reactors used 100 ml of fecal inoculate resulting in an overall volume of 400ml. The media feed was started 24 hours after inoculation at a constant rate of 400 ml per day to emulate the 24-hour average passage time in the human gut. Media was changed 16 times throughout the course of the experiment, each time media was prepared fresh. On day 13 it was discovered that the feed line to vessels 1 and 2 was blocked, this blockage could have occurred any time after day 11.

In addition to media feed rate, Oxygen, pH, temperature, and stir rate was controlled by the IRIS software (v6, Infors). Oxygen concentration in the vessels was kept below 1% via positive Nitrogen pressure at 1 LPM. Oxygen concentration was measured continuously using Hamilton VisiFerm DO Arc 225 probes. The oxygen probes were calibrated using a two-point calibration performed with Nitrogen flowing at 1 LPM as the zero-point calibration and room air flowing at 1 LPM as the 100% calibration point. pH was maintained between 6.9 and 7.1 using a 1N HCl solution and a 1N H3PO4 solution. pH was measured continuously with Hamilton EasyFerm Plus PH ARC 225 probes. The pH probes were calibrated with a 2-point calibration with standardized pH buffers at 4.00 ± 0.1 and 10.00 ± 0.1 (BDH). Vessels were maintained at 37C via the Infors’ onboard temperature control system. Vessels were continuously stirred at 100 rpm using magnetic impeller stir-shafts.

#### Bacteroides ovatus delivery

To study community dynamics in response to changes in a single bacterial taxa, we supplemented replicate vessels 1 and 4 with 2ml of isolated *B. ovatus* at 10^10^ cells/ml (estimated by optical density) suspended in anaerobic blood heart infusion (BHI) agar, and vessels 2 and 3 with 2ml of anaerobic BHI as a control on day 23. No evidence of *B. ovatus* increase was detected via community composition. Longitudinal modeling suggest effects of adding delivery media were limited to a transient 0.5 *e.i.* shift in the balance between the families Bacteroidaceae and Porphyromonadaceae (Fig. S7).

#### Sampling

For each time-point sampling of the four replicate vessels was done as follows. Prior to sampling, sampling ports were cleared with a sterile syringe and wiped clean with ethanol. Samples were collected in the following order: Vessel 1, Vessel 2, Vessel 3, and Vessel 4. Sampling consisted of the collection of 3 ml of active artificial gut culture via sterile syringe and then immediate storage in labeled and barcoded cryovials in a −80C fridge. The full list of sampled time-points including daily, hourly, and technical replicate samples is shown in Figure S4.

#### DNA Extraction, PCR Amplification, and Sequencing

For all samples, 16S rRNA gene amplicon sequencing was performed using custom barcoded primers targeting the V4 region of the gene (71) and published protocols (71–73). All samples were randomized into seven sets of 96 for all sample preparation steps. Extractions were preformed using the MoBio PowerMag Soil DNA Isolation kit (p/n 27100-4-EP) adapted for use without robotic automation. Due to the large number of samples, samples were split randomly into two pools of 336 samples for sequencing to ensure adequate read depth per sample. The final DNA concentration for pool 1 was 35.5 ng/ul and for pool 2 was 34.5 ng/ul as assessed by Picogreen assay. Both sequencing runs were standardized to 10nM and sequenced using an Illumina MiSeq with paired end 250 bp reads using the V3 chemistry kits at the Duke Molecular Physiology Institute core facilities.

#### Identifying Sequence Variants

We used DADA2 to identify and quantify sequence variants in our dataset (74). To prepare data for denoising with DADA2, 16S rRNA primer sequences were trimmed from paired sequencing reads using Trimmomatic v0.36 without quality filtering (75). Barcodes corresponding to reads that were dropped during trimming were removed using a custom python script. Reads were demultiplexed without quality filtering using python scripts provided with Qiime v1.9 (76). Bases between positions 10 and 180 were retained for the forward reads and between positions 10 and 140 were retained for the reverse reads based on visual inspection of quality profiles. This trimming, as well as minimal quality filtering, of the demultiplexed reads was performed using the function *fastqPairedFilter* provided with the *dada2* R package (v1.1.6). Sequence variants were inferred by *dada2* independently for the forward and reverse reads of each of the two sequencing runs using error profiles learned from a random subset of 40 samples from each sequencing run. Forward and reverse reads were merged for each of the two sequencing runs. Bimeras were removed using the function *removeBimeraDenovo* with *tableMethod* set to “consensus”. Finally, the two sequencing runs were merged together into a single count table.

#### Taxonomy Assignment

Initially, taxonomy was assigned to sequence variants using a Naive Bayes classifier (77) trained using version 123 of the Silva database (78). Initial taxonomic assignments were then augmented by searching for exact nucleotide matches to the Silva database. This resulted in 96% of sequence variants being classified at the family level, 85% at the genus level, and 15% at the species level.

#### Data Preparation for Modeling

After investigating the distribution of sample sequencing depth, we chose to retain only samples with more than 5,000 read counts to remove outlying samples that may have been subject to library preparation or sequencing artifacts. This step retained 99.8% of total sequence variant counts. For computational tractability and to ensure a maximal number of retained sequence variant counts, we preformed our analysis at the family level and we retained only those families that were present with at least 3 counts in more than 90% of samples. While these filters yielded only 10 bacterial families, they represented 97.7% of total sequence variant counts.

#### Construction of Phylogenetic Sequential Binary Partition

To use the PhILR transform (22) we manually created a sequential binary partition based on the phylogenetic relationships between the bacterial families in our dataset. This manual partition was created in accord with the phylogenetic relationships between bacterial families specified in Rajilic-Stojanovic and de Vos (79). The resulting sequential binary partition is given in Figure S4.

### The MALLARD Framework

To accommodate time-series with replicate observations (as depicted in Figure 1C), we refer to samples by sample point *k* ∈ {1,…, *K*} rather than time point *t* ∈{1,…, *T*} such that *K* ≥ *T*. Further, let the function *ϕ* provide a mapping between the sample point and the time point such that *ϕ*(*k*) = *t*. As in standard time-series notation, we assume that these sample point indices are temporally ordered such that *ϕ*(*k*) ≤ *ϕ*(*k* + 1) for all *k*. With this notation, denote a typical longitudinal microbiome dataset as the array ***Y*** with element *y_krd_* representing the number of counts measured for taxa *d* ∈ {1, …, *D*} in individual (or artificial gut) *r* ∈ {1, …, *R*} at sample point *k*. We also denote the total sequence counts (sequencing-depth) attributed to the *k*-th sample from the *r*-th individual as 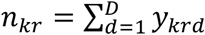.

#### Likelihood Model

High-throughput DNA sequencing does not measure the total number of target DNA transcripts in a biological system but only a random subset of this total. The size of this subset is represented by the sequencing depth of a sample. This feature of DNA sequencing leads to a competition to be counted between transcripts in which more abundant transcripts can exclude observations of less abundant transcripts. To capture this behavior, we modeled DNA sequencing as a multinomial counting process

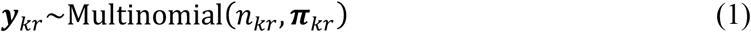

where ***π**_kr_* = (*π_krl_*, …, *π_krD_*) denotes a vector of relative abundances of taxa in a sample such that Σ*_d_ π_krd_* = 1.

While the multinomial component of our model accounted for uncertainty due to the counting process underlying DNA sequencing, the multinomial component alone was insufficient to account for the other sources of technical and biological variation in longitudinal microbiome studies. To allow for this type of extra-multinomial variability we modeled the multinomial parameters (***π***) as distributed logistic-normal. We chose the logistic-normal distribution for two reasons. First, the logistic-normal has greater flexibility than the more common Dirichlet distribution allowing for both positive and negative covariation between taxa (40, 80). Second, the logistic-normal distribution represents the central limiting distribution over the space of multinomial parameters assuming multiplicative errors (37, 81). We chose to model with a multiplicative error structure due to the multiplicative nature of bacterial growth and DNA amplification.

While the logistic-normal can be difficult to work with in terms of relative abundances, under the Isometric Log-Ratio transform (ILR) (22, 82), the logistic-normal simplifies to a multivariate normal distribution allowing many standard statistical tools to be used (37). In particular, to describe the variation of the multinomial parameters in our system we made use of a class of linear Gaussian state-space models often referred to as dynamic linear models (38). We could then write our model for the multinomial parameters ***π**_kr_* as

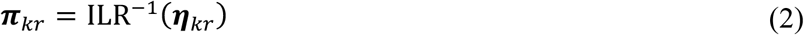

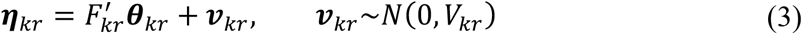

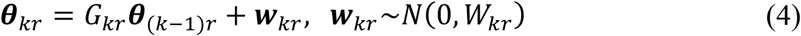

where ***η**_kr_* is the (*D* – 1)-vector representing the relative abundances ***π**_kr_* in the ILR space, ***θ**_kr_* is a *p*-vector denoting the system state, *F_kr_* is a *p* × (*D* – 1) matrix describing how the state parameters translate into observations, ***v**_kr_* is a (*D* – 1)-vector representing the measurement noise, *G_tr_* is a *p* × *p* matrix describing the dynamics of the state vector, and ***w**_kr_* is a *p*-vector representing random perturbations that influence the system state. Taken together Equations (1–4) define the likelihood component of the multinomial-logistic normal generalized dynamic linear modeling (MALLARD) framework. This likelihood model is a special case of the multinomial-conditionally Gaussian dynamic linear models of Cargnoni, Muller and West (34) where the ILR is specified as the link function and the possibility of replicate observations is made explicit by writing these models in-terms of the temporally ordered sample points rather than the time points of samples. This MALLARD framework also differs from that of Cargnoni, Muller and West (34) in terms of our prior specifications and inference methods (*see below*). Modeling flexibility in the specification of *F_t_*, *G_t_*, *w_t_*, and *V_t_*, allows the MALLARD framework to encompass many different types of models as special cases including dynamic and static mixed-effects or factor models (38). A thorough review of the use of dynamic linear models, and MALLARD models by extension, is given in West and Harrison (38).

The model used in this work is a specific MALLARD model in which the state parameters ***θ**_kr_* are identified with the unobserved microbial composition before the addition of confounding technical variation. With this specification, the dimension of the state space *p* is equal to *D* – 1. As our primary interest was in retrospective inference we chose to model the state evolution as a simple random walk (or constant level) such that 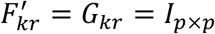, where *I_p × p_* denotes the *p*-dimensional identity matrix. As we randomized our samples through all steps of preprocessing, we assumed that the technical noise profile of each sample is identical such that we could write *V_kr_* = *V*. Additionally, we assumed that the biological variation profile is identical between each time-point such that we can write

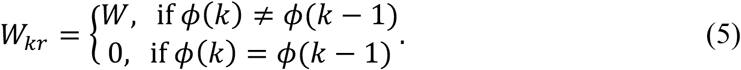

The conditional relationship in Equation (5) allowed us to account for the lack of temporal evolution between replicate samples. Finally, we chose the Phylogenetic Basis defined in Silverman, Washburne, Mukherjee and David (22), without tip or branch weights as a default basis for all posterior computations.

In this work, we chose to distinguish two types of missing observations based on whether they are to be imputed during posterior sampling. To ensure each of the four replicate vessel time-series remains synchronized we considered any sample point that is missing from one vessel but observed in at least one other vessel as a missing value to be imputed by augmenting posterior simulations with a corresponding vector ***η**_kr_* (see below). Conversely, we considered sample points that are present in none of the four replicate vessels as missing values that were marginalized over using the Kalman filter (see below) (83). We padded periods of daily sampling with missing values so that the entire dataset could be analyzed at an hourly base interval.

#### Prior Specifications

We specified two types of prior beliefs for parameters in the MALLARD likelihood model, the priors over the state vectors ***θ*** and the priors over the covariance components *V* and *W*. For the state vector component we specified a distribution over the true composition of each of the replicate vessels one-hour prior to the first observed sample such that for *r* ∈ {1,…, 4}

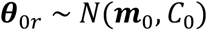

with ***m**_0_* = **o***_p_* and *C_0_* = 25 · *I_p × p_*. With this specification, there is an approximately 66% probability (1 standard deviation) that no single taxa was greater than approximately 200 times more abundant than the geometric mean of the remaining taxa. As we are only modeling bacterial families that are present with at least 3 counts in at least 90% of samples, we believed that such concentration of our prior about zero was warranted.

To quantify our uncertainty in the covariance components of our model we also specified a prior distribution for *V* and *W*. While this is most commonly done using Inverse Wishart distributions due to their conjugacy with the multivariate normal distribution (38), here we chose to use a reparameterization of the covariance matricies *V* and *W* with non-conjugate priors to improve numerical stability. A covariance matrix *Σ* ∈ {*V, W*} can be parameterized as

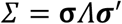

where *Λ* denotes the correlation matrix corresponding to *Σ*, and ***σ*** represents the diagonal matrix with positive diagonal entries (*σ_1_, …, σ_D_* – _1_) which dictate the scale of the covariance matrix. Thus, for the covariance matrices *V* and *W* we specified the components 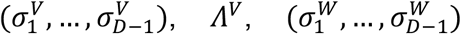, and *Λ^W^* respectively. As the representation of *Λ^V^* and *Λ^W^* depends on the chosen ILR basis and we had no prior knowledge regarding how the technical and biological variation will decompose in our chosen basis, we chose to use a uniform prior over the space of symmetric positive semi-definite matrices such that

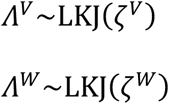

with *ζ^V^* = *ζ^W^* = 1 (84). With regards to the terms 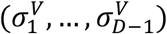 and 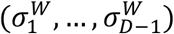, we chose independent log-normal priors to ensure that these terms were strictly positive and to allow parameterization of uncertainty with respect to multiplicative fold-changes. In particular, for each *i* ∈ {1, …, *D* - 1} we specified

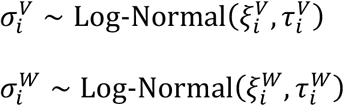

with 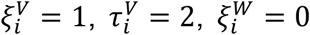, and 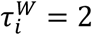. This specification reflects a 95% probability that the ratio of total technical 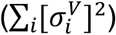 to total biological variation 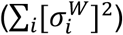 is between 10^-2^ and 10^2^ with an expected value of approximately 10.

#### Posterior Inference

Prior work with multinomial conditionally-Gaussian dynamic linear models used Metropolis-within-Gibbs sampling schemes (34). However, our preliminary results suggested that mixing rates under such schemes were too slow to be practically useful for microbiome data. Instead, we chose to use Hamiltonian Markov Chain Monte Carlo (HMCMC), for all posterior simulations. To reduce the dimension of Markov chain parameter space we exploited the Kalman filter to marginalize over all state parameters (***Θ***) (38, 85).

Before describing the inference method in detail, we introduce the following notations, for ϱ ∈ (1, …, *p*) let ***H*** = {*η_krϱ_*} and let ***Θ*** = {*θ_krϱ_*} represent the collection of all ILR transformed multinomial parameters and all state parameters respectively. Our goal was to obtain samples from the posterior distribution *P*(***Θ**, **H**, V, W* | ***Y***). As this distribution may be factored as *P*(***Θ**, **H**, V, W* | ***Y***) = *P*(***Θ** | ***Y**, **H**, V, W**)*P*(***H**, V, W* | ***Y***), we could first generate samples from *P*(***H**, V, W* | ***Y***) by HMCMC and then conditioned on these samples use the Kalman filter and smoother to generate samples from *P*(***Θ** | ***Y**, **H**, V, W**)(86, 87). This approach essentially removed *K* × *p* × *R* parameters from HMCMC and instead sampled these parameters directly from their posterior distribution using the closed form updates provided by the Kalman filter. The posterior distribution *P*(***H**, V, W* | ***Y***) may be written as

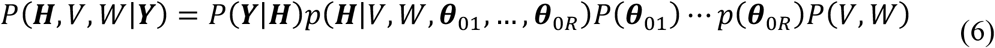

The term *P*(***Y*** |***H***) is the multinomial likelihood for the observed data given the multinomial parameters such that 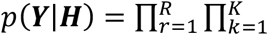 Multinomial(*n_kr_*, ILR^-1^(***η**_kr_*)). Using the first order Markov structure of the MALLARD likelihood model, the middle terms in Equation (6) can be expanded to

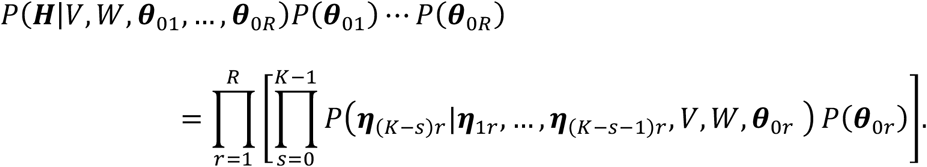

The terms within the square brackets correspond to the 1-step ahead predictive densities that can be calculated directly from the Kalman filter (38), thus requiring *R* independent Kalman filter calculations to evaluate this contribution to the posterior density. Finally, the term *P*(*V,W*) represents

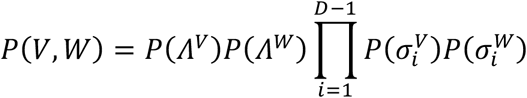

each term of which can be calculated directly from their defined prior distributions.

Posterior inference was performed using the No-U-Turn Sampler (NUTS) provided in the Stan modeling language (88, 89) using 4 chains run in parallel each with 1000 transitions for warmup and adaptation and 1000 iterations collected as posterior samples. Preliminary results suggested that the time required for the NUTS sampler to converge to the typical set could be quite sensitive to entirely random parameter initializations. To address this computational limitation, the following parameters were manually specified: ***H*** was set by first adding a pseudo-count of 0.65 to each observed counts and then normalizing the counts of each sample to sum to 1, *Λ^V^* and *Λ^W^* were each initialized to the *p × p* dimensional identity matrix, all other parameters were randomly initialized. To mitigate the potential bias introduced by fixing these parameters during initialization, approximate posterior samples from the model were first drawn using a variational algorithm (90) and then 4 randomly selecting posterior samples from this approximate posterior sample were used to initialize 4 parallel HMCMC chains. Convergence of the chains was determined both by manual inspection of sampler trace plots and through calculation of the Gelman-Rubin statistic (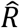) (91). All sampled parameters had an 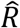 value less than 1.01. Posterior intervals for all calculations derived from directly sampled quantities were calculated by preforming the necessary computations on each posterior sample independently and then summarizing the resulting distribution over calculated quantities.

### Comparing the Correlation Structure of Technical and Biological Variation

To quantitatively compare the correlation structure of technical and biological variation we analyzed the probability that posterior samples of the correlation matrices corresponding to *V* and *W* came from the same distribution using a permutation scheme and a distance metric on the space of square symmetric positive semi-definite matricies. To isolate our inferences to only involve the correlation structure of *V* and *W* and not the magnitude of the variation, we transformed sampled covariance matrices into corresponding correlation matrices which we denote *V^c^* and *W^c^*. We took as a measure of distance between two correlation matrices *S_1_* and *S_2_* the Riemannian metric on the space of square symmetric positive definite matrices defined by

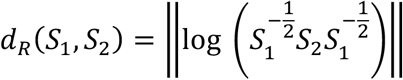

as described in (92) and calculated using the function *distcov* with option *“Riemannian”*in the R package *shapes* (93). Using this distance metric, we calculated a distance matrix between 500 posterior samples of *V^c^* and *W^c^* each. Let *D* represent the resultant 1000 by 1000 distance matrix such that *D_ij_* represents the distance between correlation matrix *i* and correlation matrix *j*. Let *l* represent the vector of labels of the correlation matrices such that *l_i_* ∈ {*V^c^,W^c^*} for *i* ∈ {1, …, 1000}. We define a statistic *δ* as the ratio of the within to between group distances in *D*

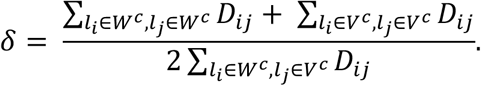

Thus, low values of *δ* indicate that most of the distance between the correlation matrices is attributable to intergroup differences in correlation structure whereas larger values suggest that most of the pairwise distances come from differences within groups. We built a distribution of *δ* under a model in which the sampled correlation matrices all came from the same distribution by permuting the labels *l* and recomputing *δ* 1000 times. The probability that posterior samples of *V^c^* and *W^c^* came from the same distribution was calculated by calculating the probability of a test statistic more extreme than that which we observed under the created permutation distribution.

### Computation of Total Technical and Biological Variation

Denoting the trace of a matrix as *Tr*(·), Total biological and total technical variation were calculated as *Tr*(*W*) and *Tr*(*V*) respectively. The proportion of variation attributable to biological sources was calculated as *Tr*(*W*)/(*Tr*(*V*) + *Tr*(*W*)). Based on the linear systems assumption underlying our model, we choose to calculate the total biological variation as a function of sampling interval *L* as *L* · *Tr*(*W*) (Fig. 3B). Technical variation does not vary with sampling interval and was therefore held constant as a function of sampling interval.

### Changing Representations of Posterior State Estimates and Covariance Matrices

While we chose the PhILR basis (22) for all posterior computations as well as initial analysis (Fig. 4 and S6,7), we also made use of the isometric properties of the ILR transform to represent our posterior state estimates and our posterior samples of covariance matrices *W* and *V* with respect to alternative coordinates. In what follows we denote vectors or matrices represented with respect to something other than the PhILR coordinates with an associated superscript. We used the inverse PhILR transform to represent any vector ***x***= (*x_1_*, …,*x_D_*_-1_) in terms of raw relative abundances such that ***x^*^*** = PhILR^-1^ (***x***) (22). Any other log-ratio quantities could then be calculated from ***x^*^***. Transforming a covariance matrix *Σ* between representations was done by making use of the following identities (37)

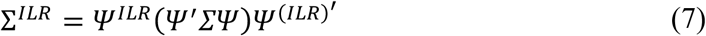

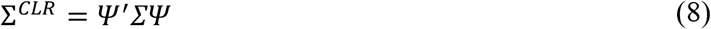

where *Ψ* denotes the contrast matrix of the PhILR transform (22) and *Ψ^ILR^* represents the contrast matrix of an arbitrary ILR transform. In addition, we made use of the variation matrix representation of a covariance matrix in which the covariance matrix is represented as a matrix describing the variance of all pairwise log-ratios (40). Letting *ρ_ij_* represent the entry in the *i*-th row and *j*-th column of *Σ^CLR^*, we compute the variation matrix *T* element-wise as (37)

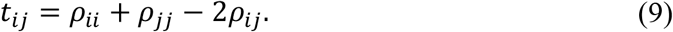

### Hierarchical Clustering of Variation Matrix

To develop a sparse representation of the principle directions of biological variation in our dataset, we make use of an algorithm for determining a sequence of orthonormal balances that maximize successively the explained variance in a dataset (principle balances) (94). As *T^W^*, the variation matrix calculated from *W*, is proportional to the Aitchison distance between the bacterial families in our dataset (94, 95) applying Ward clustering to the matrix *T^W^* results in an approximate solution to the problem of determining principle balances (94, 96). For each posterior sample of *W* we calculated *T^W^* using Equations (8) and (9). We then calculated a sequential binary partition from each posterior sample of *T^W^* using Ward clustering (97). To summarize this posterior sample of sequential binary partitions we computed the majority-rule consensus tree and the frequency with which a given bipartition occurred in our posterior sample using the functions *consensus* and *prop.part from* the R package *ape* (98).

### Exploring the Relation Between Starting Composition and Biological Variation

To investigate the relationship between starting community composition and biological variation in our replicate vessels we made use of the CLR representation of both the community state at the first observed time-point (***θ***_1*r*_) and the biological variation (*W*). The mean composition in the first observed sample 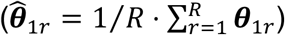 represented in the CLR basis was used as a measurement of the starting community. As we were primarily interested in the relative degree to which bacterial families vary in our dataset, we normalized the diagonal elements of the CLR representation of the biological variation matrix *W* to sum to 1 forming a composition and then used the CLR transform of these normalized variances as our measure of the relative biological variation of each bacterial family. We represent the resulting CLR transformed relative biological variation of each bacterial family by the vector *ω* = (*ω, …, ω_D_*). For each posterior sample, a univariate linear regression model given by the relation 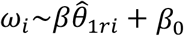 where *β* denotes the slope of fitted model and *β_0_* represents an intercept was fit resulting in a posterior sample over *β*. Probability contours over the joint posterior distribution of each pair 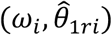 was calculated using kernel density estimates computed by the R package *ks* (99).

We constructed a permutation distribution to investigate whether the negative relationship between 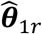 and ***ω*** suggested by this posterior distribution of *β* could be trivially due to there being more low abundance families than high abundance families in our starting community or a feature of working with such CLR transformed variables. This permutation distribution was constructed by randomly permuting the labels of the vector 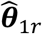 and re-computing the posterior mean of *β* 1000 times. We computed the probability of our observed posterior mean of *β* under the constructed permutation distribution to estimate the probability, conditioned on our prior beliefs, that the observed inverse relationship was simply due to the rank abundance distribution of our starting community or a feature of working with such CLR transformed variables. This probability is different than a traditional frequentist p-value in that it is conditioned on our prior specifications and our observed data.

### Sensitivity Analyses

The sensitivity of our results to our prior specifications was assessed by rerunning our posterior inference and computations under perturbed priors. We identified two key quantities in our analysis, the regression slope *β* and the percentage of total variation attributable to biological sources *Tr*(*W*), and looked at the change of the associated posterior intervals under the perturbed priors. The results and specifications of the modified priors are shown in Figures S5 and S12.

### Calculation of Fold Changes from Balance Values

Changes in balances values are a measure of evidence information (39). Given a composition ***x*** with *D* taxa we can denote the balance that separates a group of *r* taxa from another group of *s* as

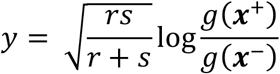

where we denote the geometric mean of the elements of ***x*** that correspond to each of the two groups as *g*(***x***^+^) and *g*(***x***^−^) respectively. In this form, the equivalent fold change between geometric means of the two groups can be calculated as

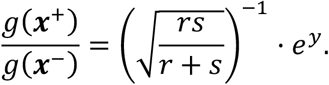

### Data Simulation

To test our implementation and explore the behavior of the MALLARD model we used to analyze our artificial gut dataset, we simulated a toy microbial community time-series of three bacterial taxa (Fig. S15 A,B) based on the likelihood component of this model. We simulated a single vessel over 200 time-points with an extra 25 replicate samples taken on the final time-point of the series. We also modeled missing observations (n=3), sparsity (23% of counts were zeros), and many low counts (53% of counts were less than or equal to 10). We created a random ILR basis denoted by the contrast matrix ***ψ**^true^* by using the functions *named_rtree, phylo2sbp*, and *buildilrBasep* from the R package *philr* (100). With respect to ***ψ**^true^*, we simulated data in accordance with our likelihood model using the following “true” values,

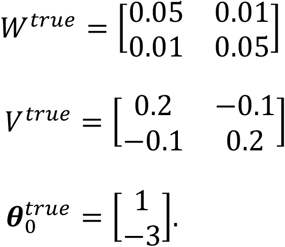

After simulation, we removed samples from time points 15, 16, and 20 to simulate missing observations. To demonstrate that our inferences were not sensitive to our choice of basis, we modeled the resulting dataset using a second random ILR basis ***Ψ***. Our results in comparison to the true values are shown in Figure S15 with respect to the basis ***Ψ***.

Analysis of this simulated dataset showed that estimates for the unobserved compositions η and θ were closer to the true simulated values than a standard modeling approach of normalizing read counts to proportions (Fig. S15 C,D). This result is clearest in regimes rich in low and zero read counts, which we expected because of our Bayesian approach to modeling the read counting process. In addition, we found that the model correctly estimated distinct technical and biological variation patterns (Fig. S15 E,F). Thus, our simulations suggested that our model implementation could successfully decompose longitudinal microbiota data into component processes and characterize technical and biological variation.

### Visualizations

All visualization were produced using the R programming language (v3.4.0) with the addition of the following packages: *ggplot2* (101), ggtree for plotting phylogenetic trees (102), and *compositions* (103).

### Code and Data Availability

Demultiplexed sequencing data were submitted to the NCBI SRA database under accession number SRP140877. All code has been made available at https://github.com/LAD-LAB/MALLARD-Paper-Code.

## ACKNOWLEDGEMENTS

We thank Rachel Silverman, Lionel Watkins, Firas Midani, Max Villa, Zachary Holmes, Brianna Petrone, B. Jesse Shapiro, Jonathan Friedman, and Susan Holmes for their manuscript comments. JS and LAD were supported in part by the Duke University Medical Scientist Training Program (GM007171), the Global Probiotics Council, a Searle Scholars Award, the Hartwell Foundation, an Alfred P. Sloan Research Fellowship, and NIH 1R01DK116187-01. SM would like to acknowledge the support of grants NSF IIS-1546331, NSF DMS-1418261, NSF IIS-1320357, NSF DMS-1045153, and NSF DMS1613261. This work used a high-performance computing facility partially supported by grant 2016-IDG-1013 (“HARDAC+: Reproducible HPC for Next-generation Genomics”) from the North Carolina Biotechnology Center.

## COMPETING INTERESTS

The authors declare that they have no competing financial, professional, or personal interests that might have influenced the performance or presentation of the work described in this manuscript.

**Supplemental Figure 1.**
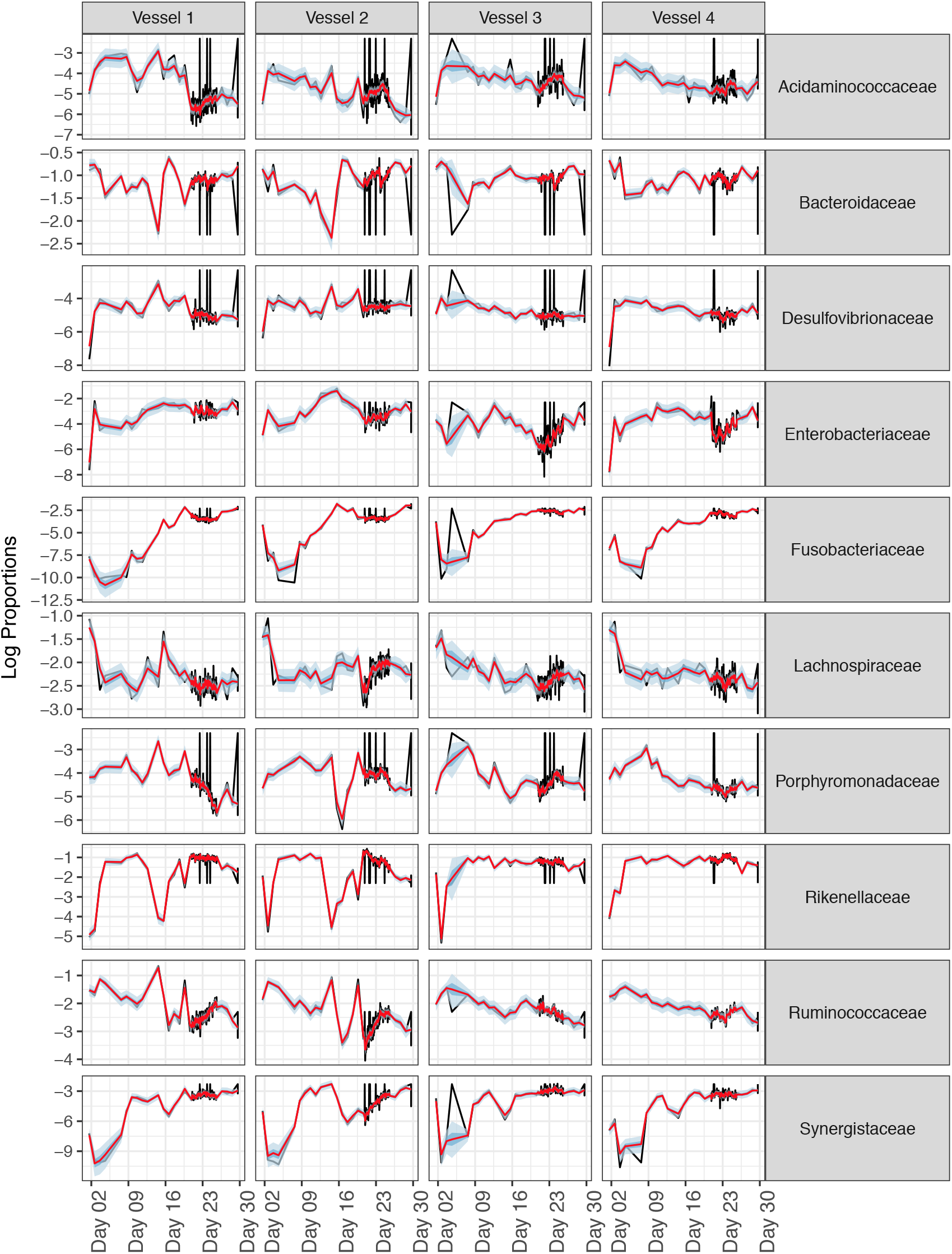
Model fits to the observed data. Posterior mean, 50% and 95% credible intervals for *θ_t_* in terms of log transformed proportions. The raw count data are shown in black for comparison. A pseudo-count of 0.65 was added to the raw data prior to normalization and log-transformation to avoid taking the log of zero values.

**Supplemental Figure 2.**
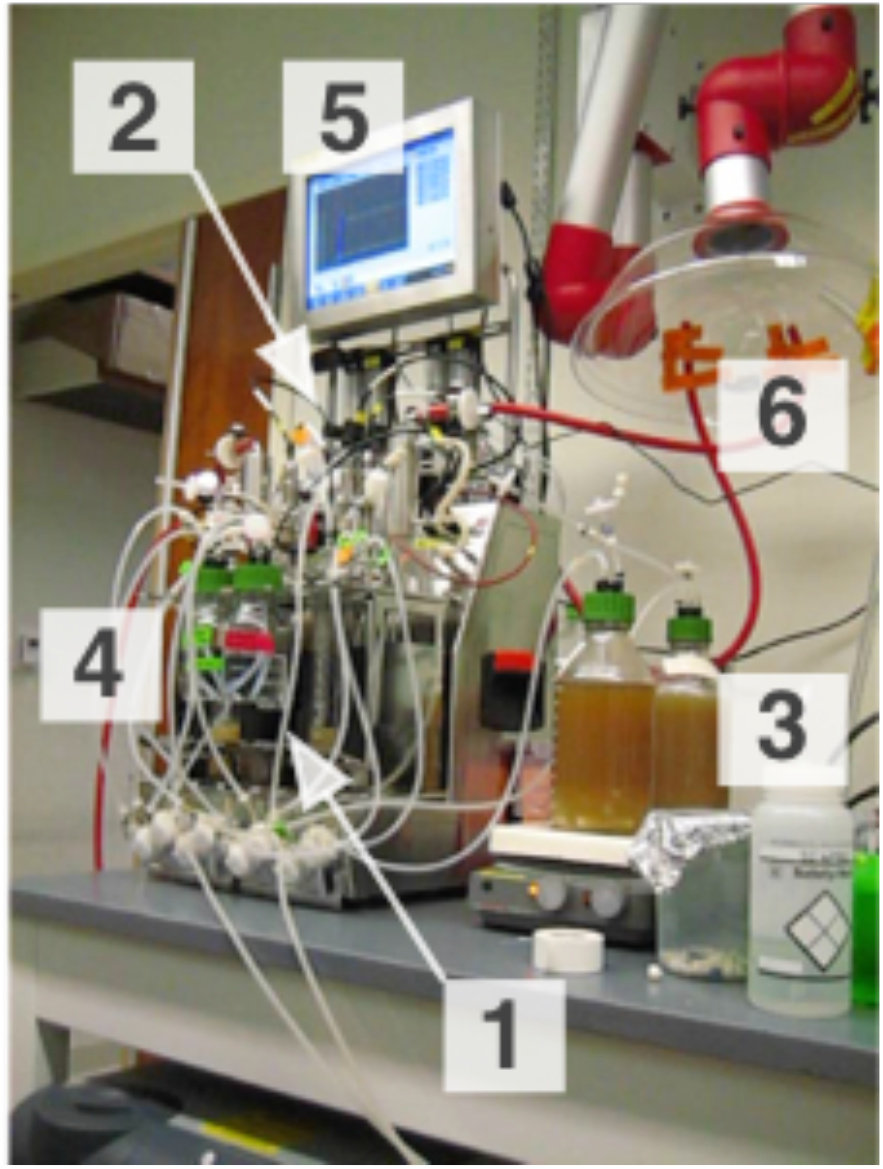
Artificial gut setup. (**1**) Reactor vessels; (**2**) flow meters controlling gas inputs; (**3**) pump-fed media; (**4**) acid and base to regulate pH; (**5**) central controller; (**6**) snorkel for exhaust. Note that only two of four replicate vessels illustrated in this photograph.

**Supplemental Figure 3.**
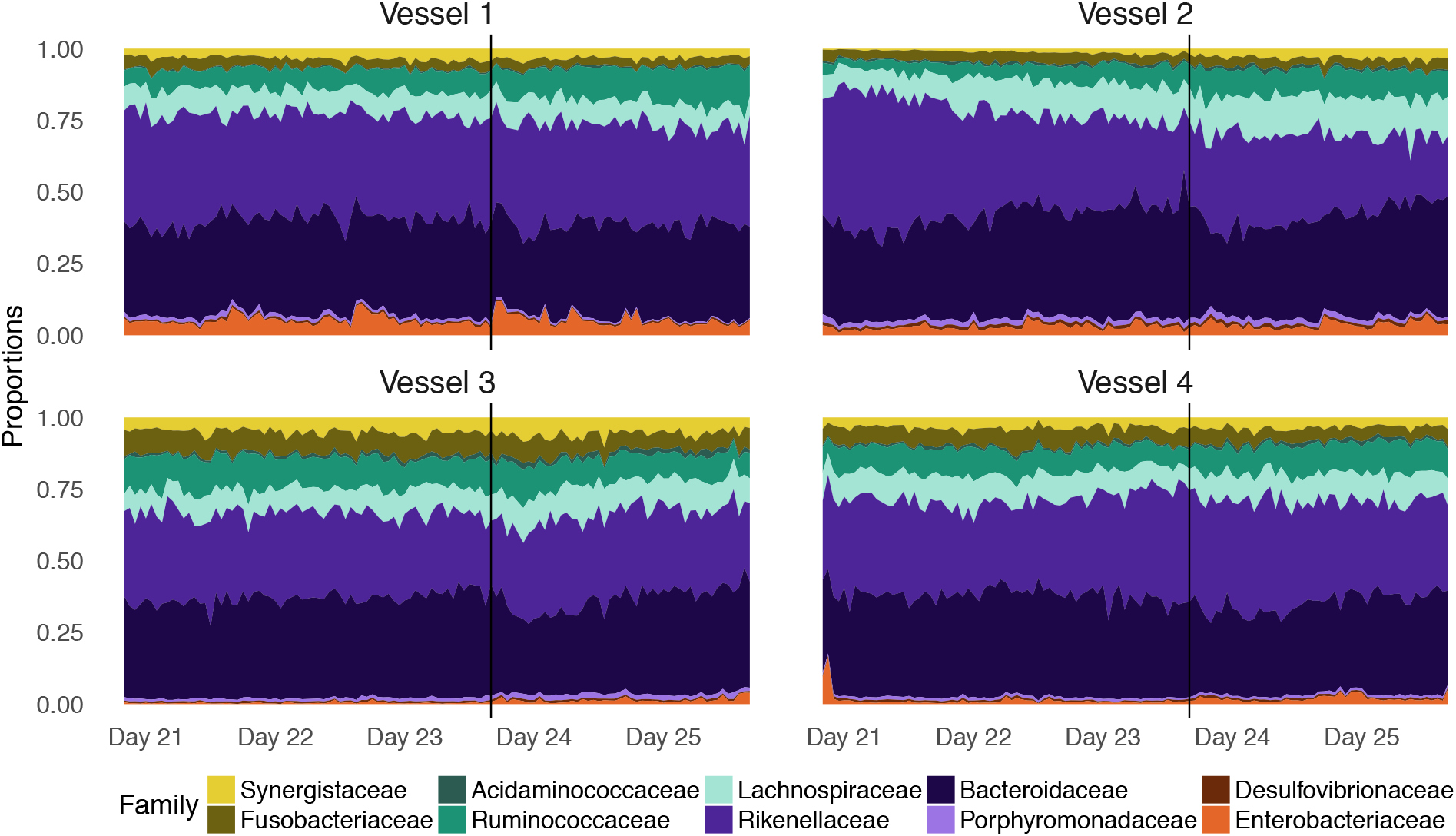
Proportions of most abundant bacterial families estimated from count data during hourly sampling period. Proportions were estimated by dividing observed counts by the total number of counts observed for each sample.

**Supplemental Figure 4.**
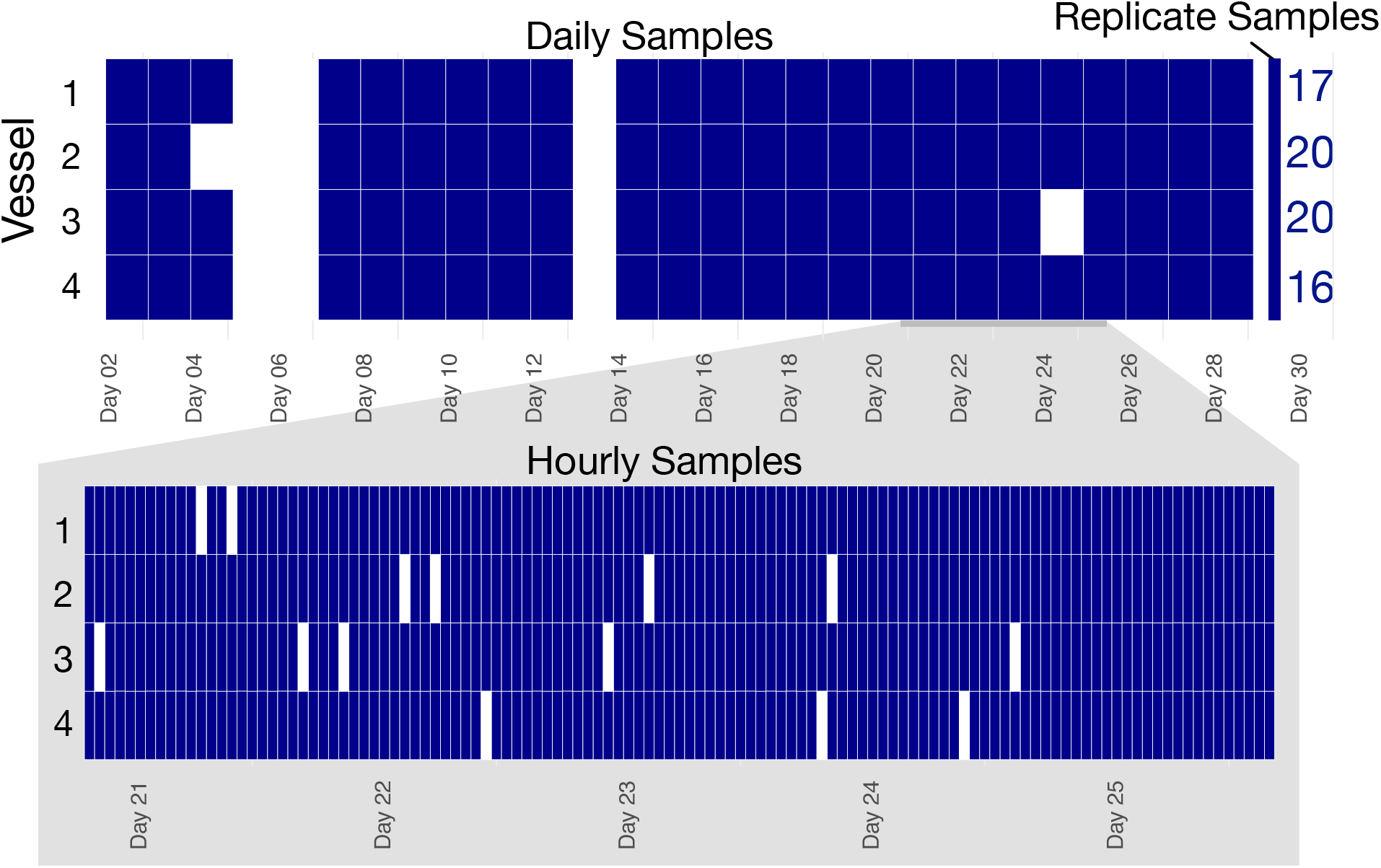
Samples were collected over a one month period with both daily and hourly sampling intervals. Daily samples were collected at 15:00±00:30 hours, hourly samples were collected within ±10 minutes of depicted time. Samples that were either not collected or filtered from analysis due to low sequencing depth are shown in white, samples that were included in analyses are shown in blue. In addition to standard hourly and daily longitudinal sampling, 20 samples were collected from the final time-point of each replicate vessel. The number of replicate samples that were included in the analysis are depicted in blue.

**Supplemental Figure 5.**
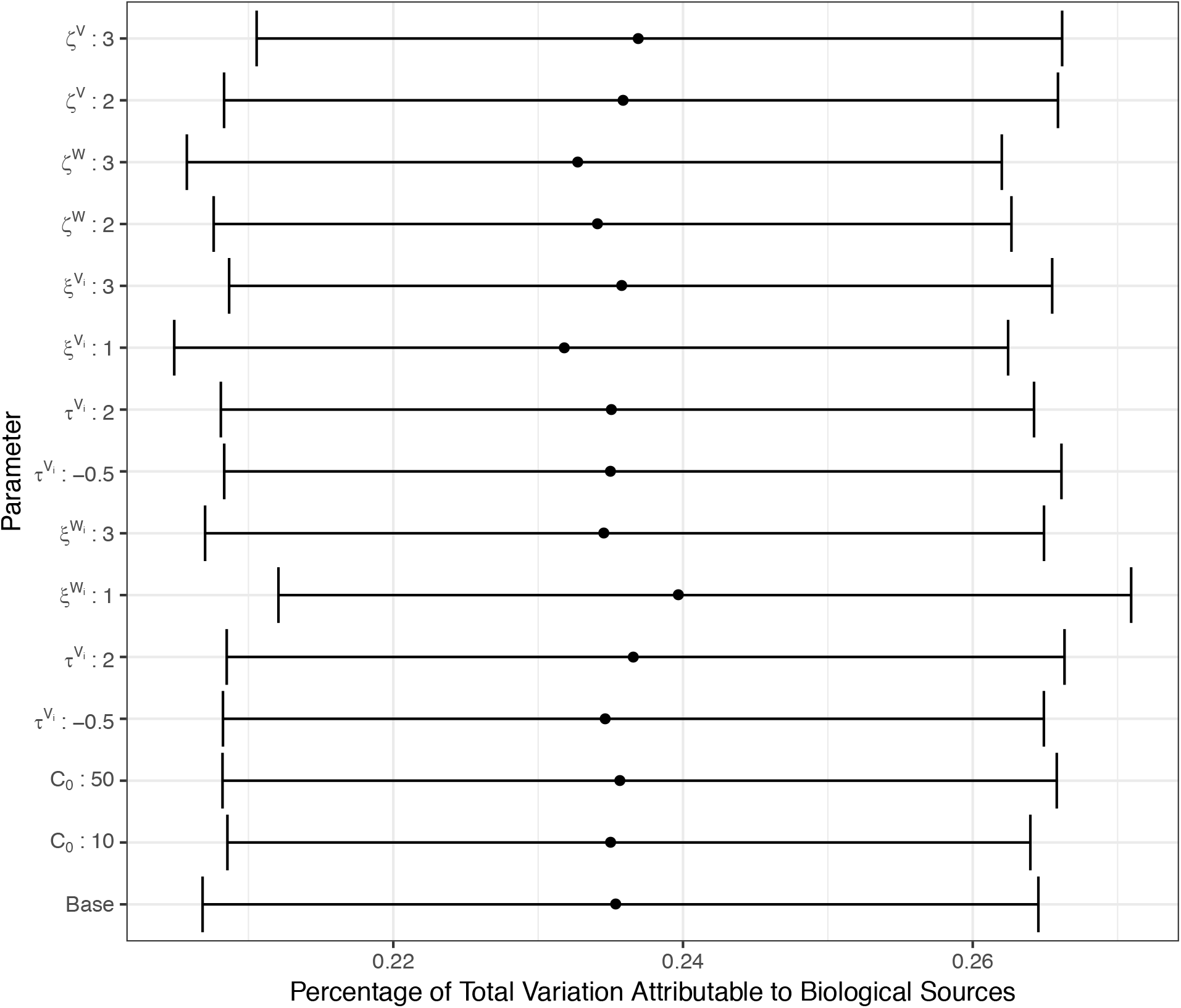
Posterior estimates for the percent of total variation attributable to biological sources is not sensitive to modification of prior parameters. The “Base” prior parameter values refer to the values specified throughout the *Methods* section. In addition, the complete model was rerun with 14 separate prior parameters settings, each deviating from the Base values with respect to one parameter. Posterior 95% credible intervals and mean are shown for each set of prior parameters.

**Supplemental Figure 6.**
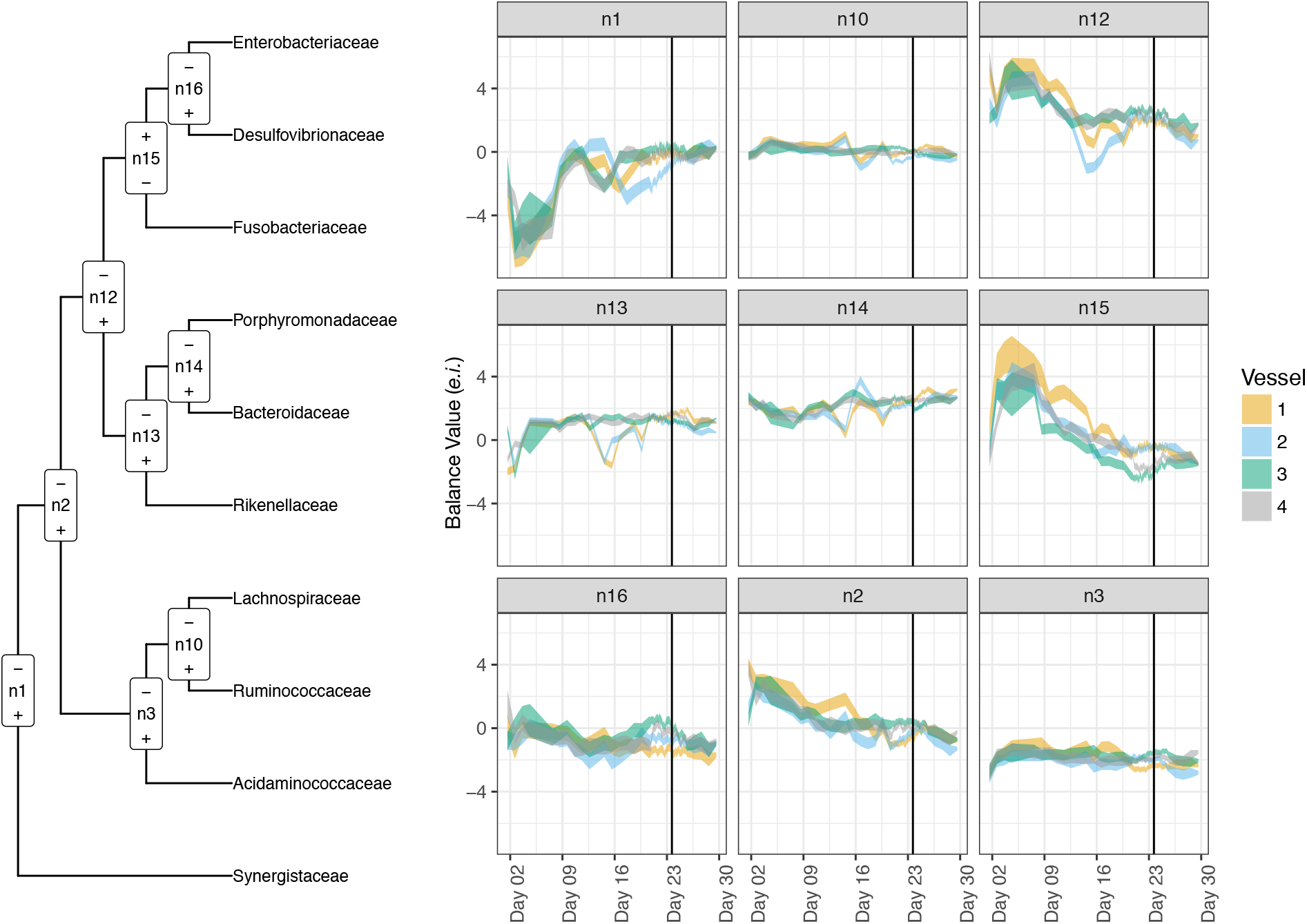
Posterior 95% credible regions for bacterial dynamics (*θ*) in the PhILR Basis. (**Left**) The hierarchical tree of phylogenetic relationships between the bacterial families with PhILR balances (n1-n16, non-consecutive numbering) depicted (*<u>Methods</u>*). Balance n12 is highlighted in Figure 3. Branch lengths are not to scale. (+) and (−) refer to which subclade is found in the numerator or denominator of the balance respectively. (**Right**) Posterior 95% credible regions for the bacterial dynamics for each PhILR balance is depicted. The time-point corresponding to *B. ovatus* treatment is depicted as a black line.

**Supplemental Figure 7.**
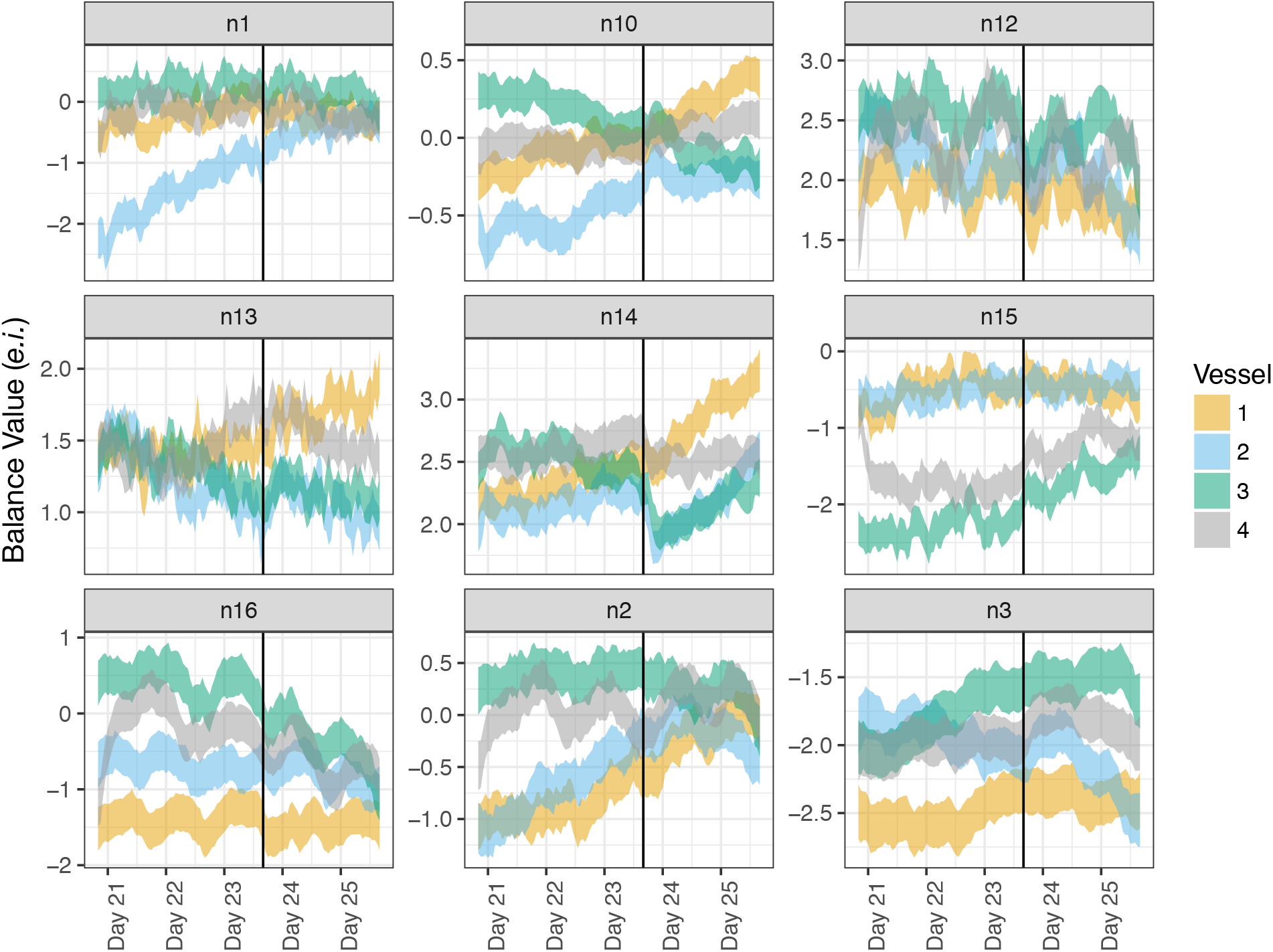
Posterior 95% credible regions for bacterial dynamics (*θ*) for hourly samples in the PhILR Basis. Balances are defined in the left panel of Figure S6, here highlighting the dynamics observed during the hourly sampling period. Balance n12 is highlighted in Figure 3. Balance n14 shows a decrease in the ratio of the families Bacteroidaceae to Porphyromonadaceae. It is likely that this effect was due to the effects of fresh delivery media as balance shifts appear strongest in the control vessels that received sham treatment (media alone) (#2 and #3) compared to the treatment vessels that received media and *B. ovatus* (#1 and #4).

**Supplemental Figure 8.**
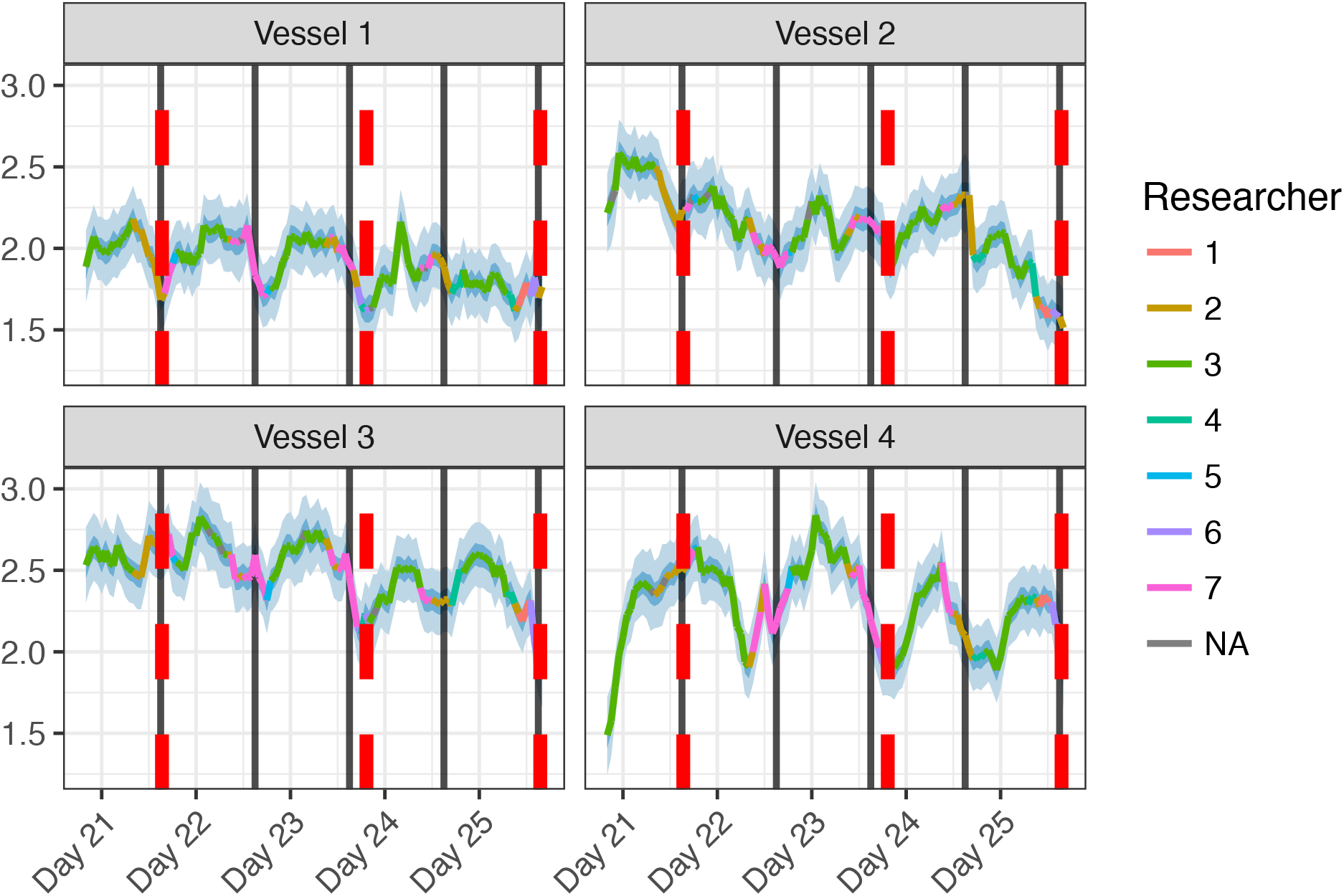
Irregular sub-daily oscillation observed in PhILR balance n12 does not correlate with known external factors. As in Figure 4B, the posterior mean and 95% credible interval of the microbial dynamics (*θ*) for balance n12 is shown during hourly sampling. The posterior mean is colored with the ID of the researcher who obtained each corresponding sample. Samples that were dropped from analysis due to low sequencing depth are denoted by NA for researcher ID. Times at which media feed bottles were changed are indicated with red dashed lines. Time-points corresponding to the daily sampling regimen are indicated by dark grey lines.

**Supplemental Figure 9.**
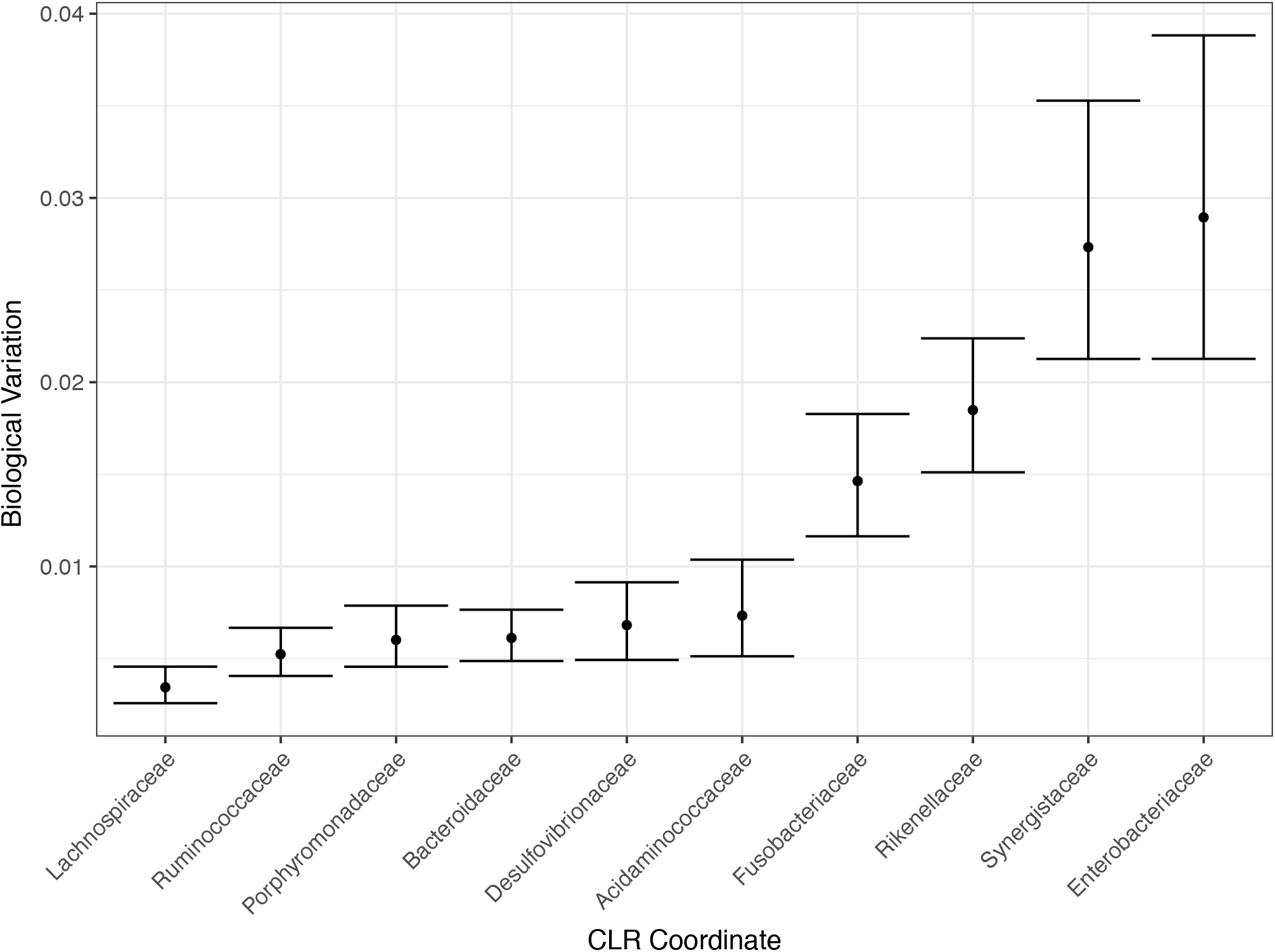
Four bacterial families comprise most the biological variation in our study. Relative biological variation for each family was taken as the corresponding diagonal entry of the CLR transformed Biological variation matrix (W^*CLR*^; *Methods*). The median and 95% posterior credible interval are depicted for each bacterial family.

**Supplemental Figure 10.**
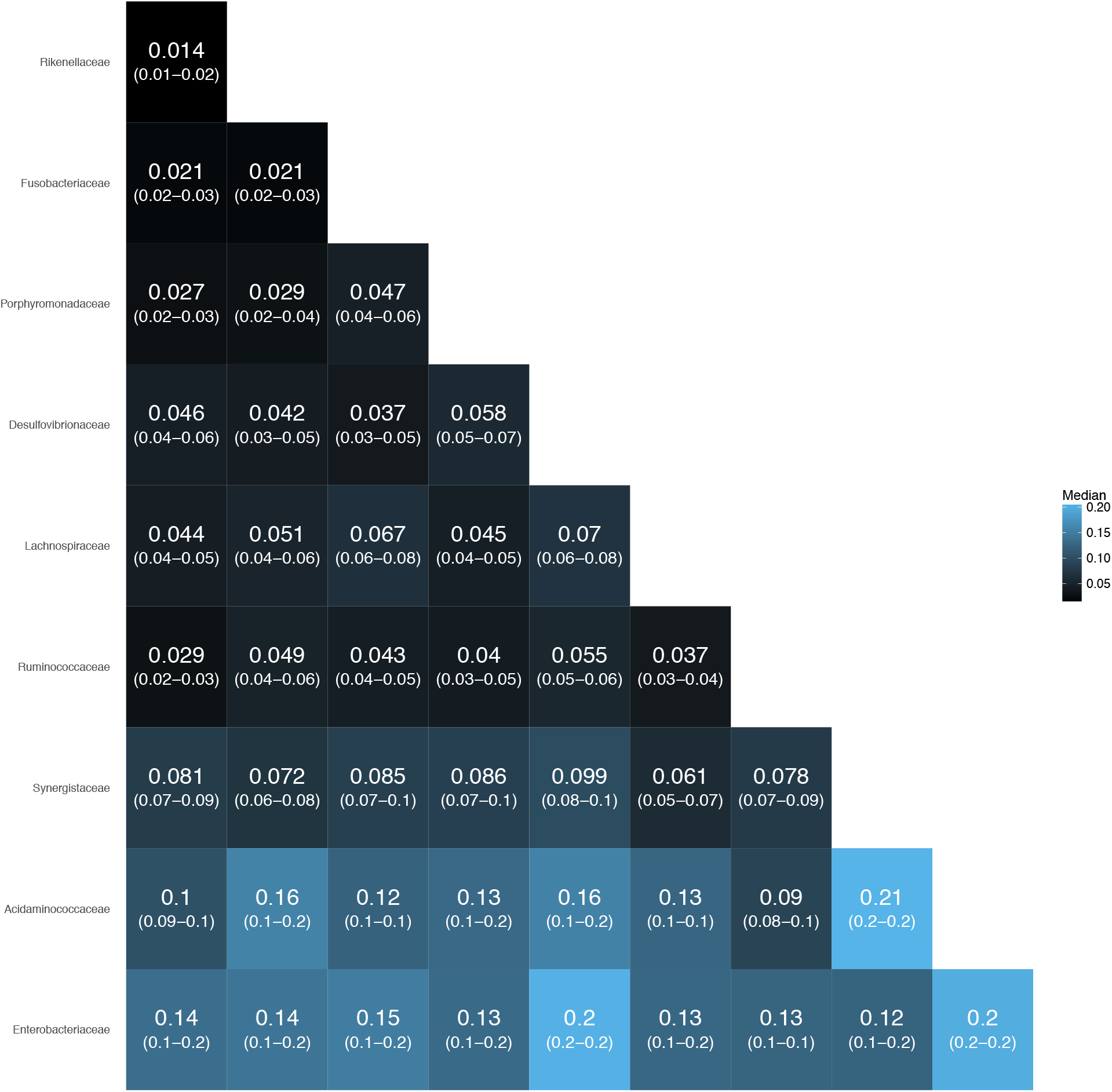
The decomposition of technical variation among bacterial families. Posterior distribution for log-ratio variance (*ρ*) between pairs of bacterial families for technical variation (*V*). Heatmap color is given by the median of the posterior distribution of *ρ*. Each cell also gives the median and 95% credible region for the log-ratio variance (*ρ*) for the corresponding bacterial families. Columns and rows refer to the bacteria in the numerator and denominator of the corresponding log-ratios respectively.

**Supplemental Figure 11.**
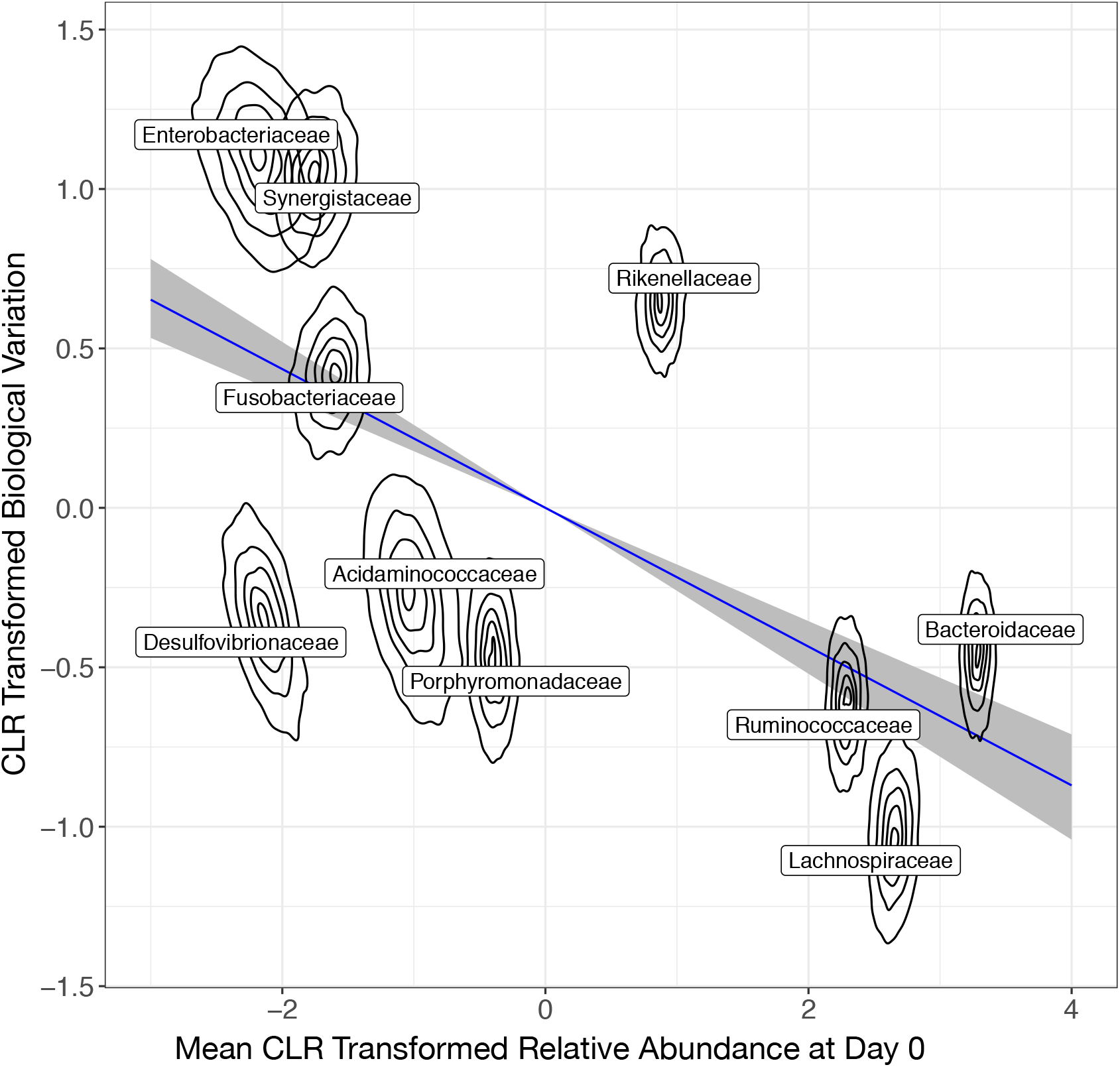
An inverse relationship between biological variation and initial relative abundance. 5, 25, 50, 75, and 95% highest posterior density regions of the posterior distribution of mean relative abundances on day 0 and biological variation of each bacterial family. Both axes are CLR-transformed. Posterior mean and 95% credible regions are also shown for the regression between these variables (*Methods*).

**Supplemental Figure 12.**
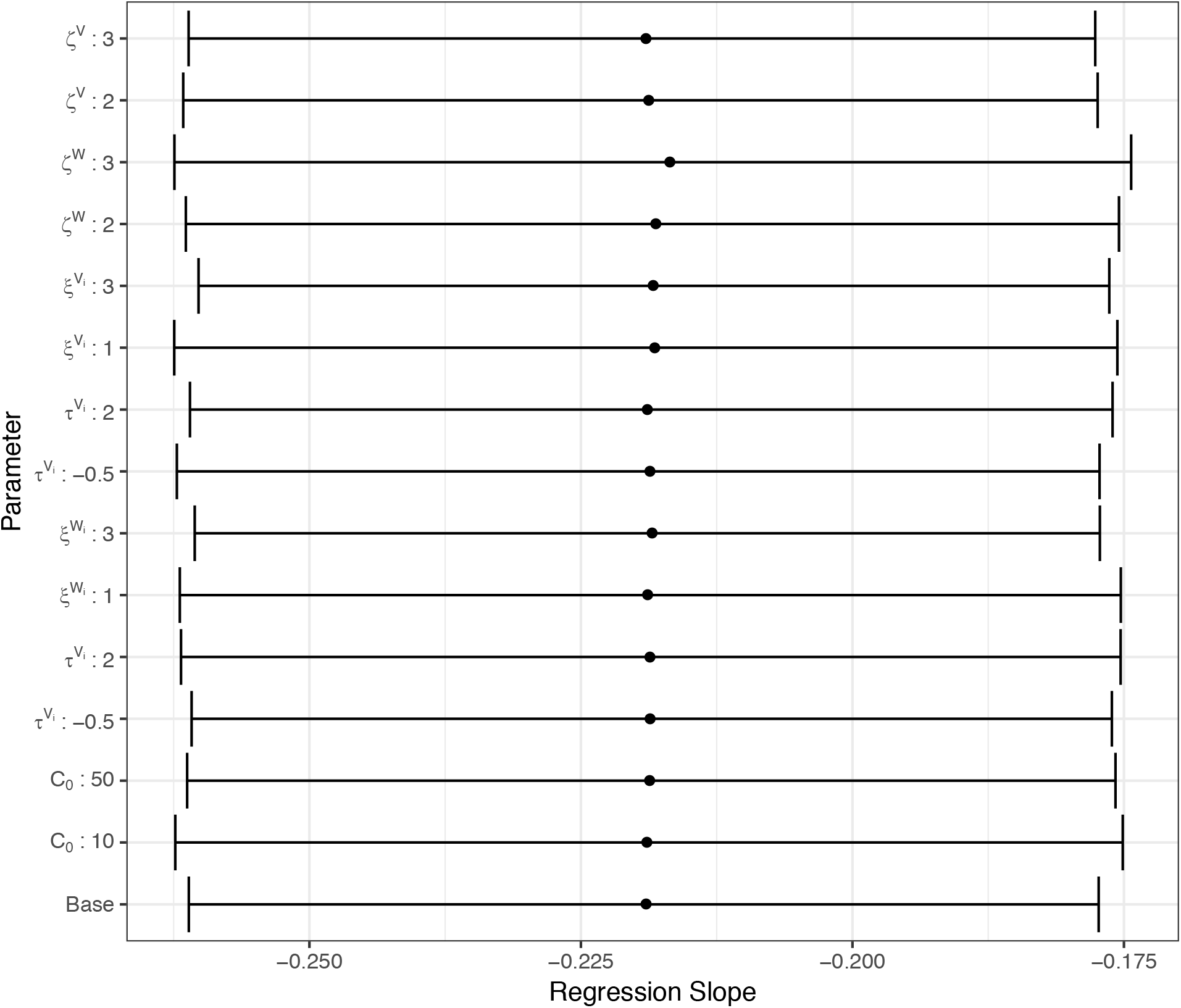
Posterior estimates for the regression slope between biological variation and bacterial family starting relative abundance is not sensitive to modification of prior parameters. The “Base” prior parameter values refer to the values specified throughout the *Methods* section. In addition, the complete model was rerun with 14 separate prior parameters settings, each deviating from the Base values with respect to one parameter. Posterior 95% credible intervals and mean are shown for each set of prior parameters.

**Supplemental Figure 13.**
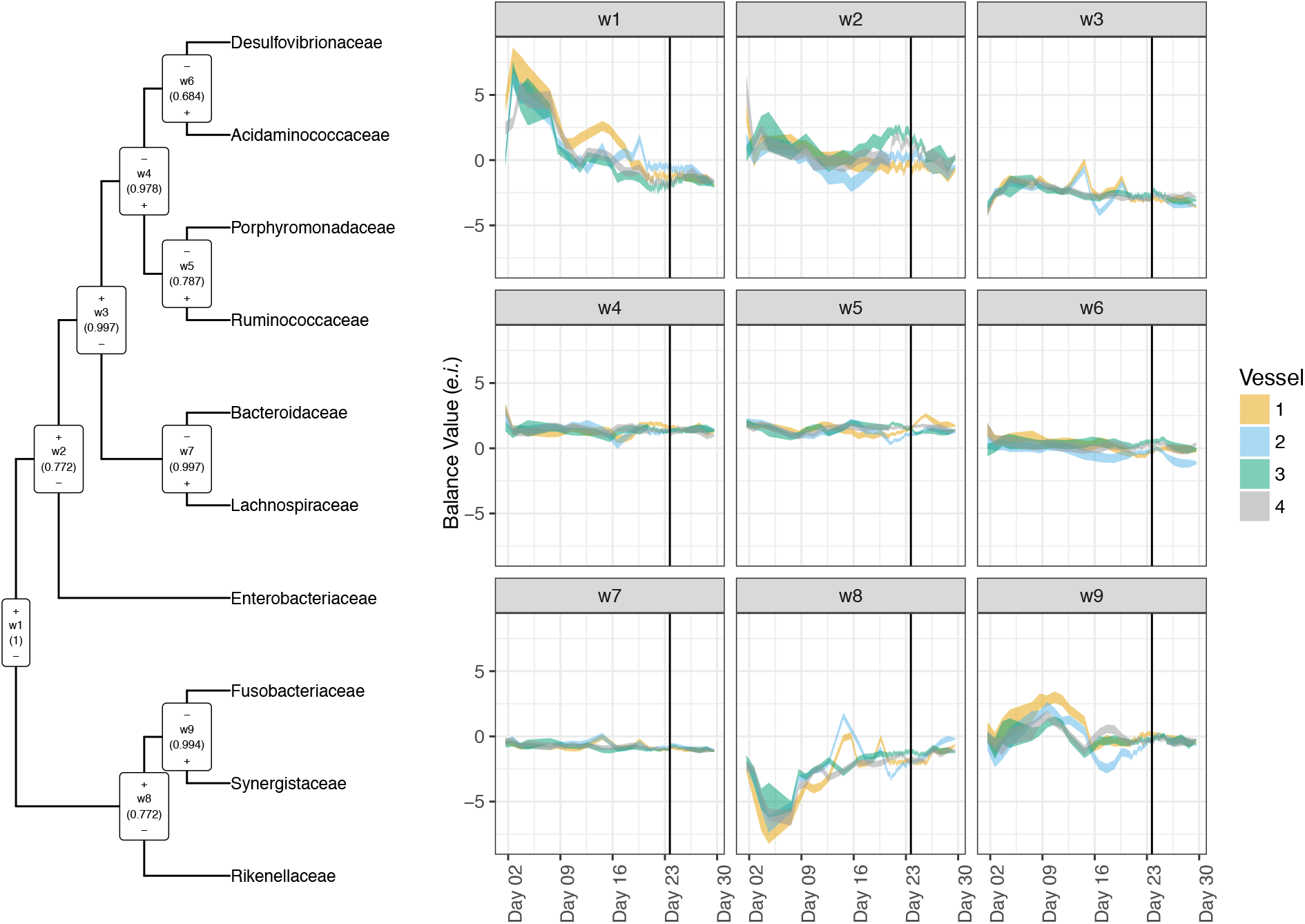
Posterior 95% credible regions for bacterial dynamics (*θ*) in the Ward Basis. (**Left**) The consensus tree created by Ward clustering of bacterial families with Ward balances (w1-w9) is depicted (*Methods*). Balances nearer the root of the tree display higher variance than balances nearer the tips (*Methods*). (+) and (−) refer to which subgroup is found in the numerator or denominator of each Ward balance respectively. The proportion of samples from the posterior distribution in which a given bipartition was present is denoted under the corresponding balance name (*Methods*). (**Right**) Posterior 95% credible regions for the bacterial dynamics for each Ward balance is depicted. The time-point corresponding to *B. ovatus* treatment is depicted as a black line.

**Supplemental Figure 14.**
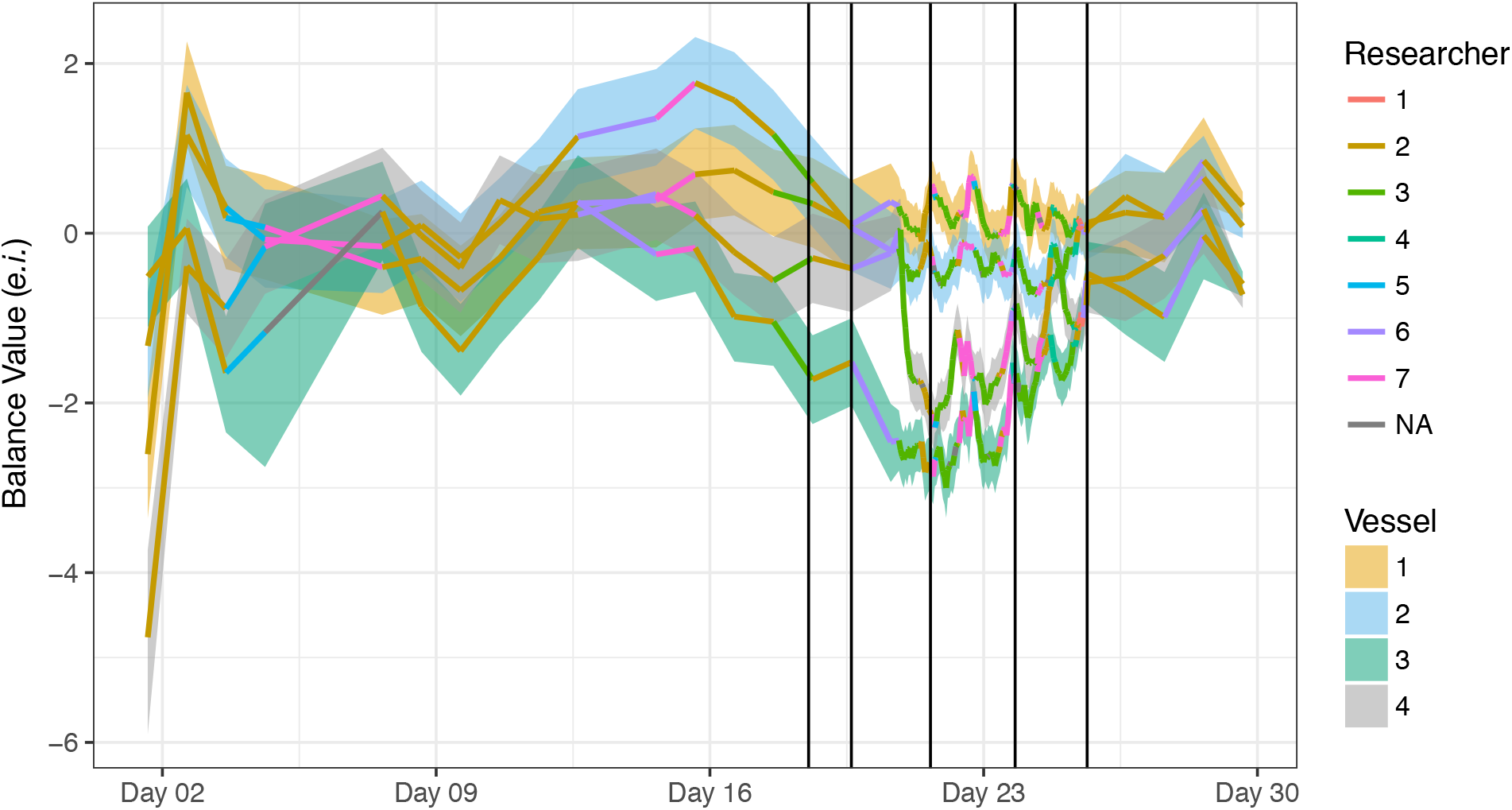
Dynamics inferred in the balance between the Enterobacteriaceae and all other taxa does not correlate with known external factors. As in Figure 5D, the posterior 95% credible interval of the microbial dynamics (*θ*) for the balance between the Enterobacteriaceae and all other taxa shown. The posterior mean is colored according to the ID of the researcher who obtained each corresponding sample. Samples that were dropped from analysis due to low sequencing depth are denoted by NA for researcher ID. Time-points corresponding to the daily sampling regimen are shown in black.

**Supplemental Figure 15.**
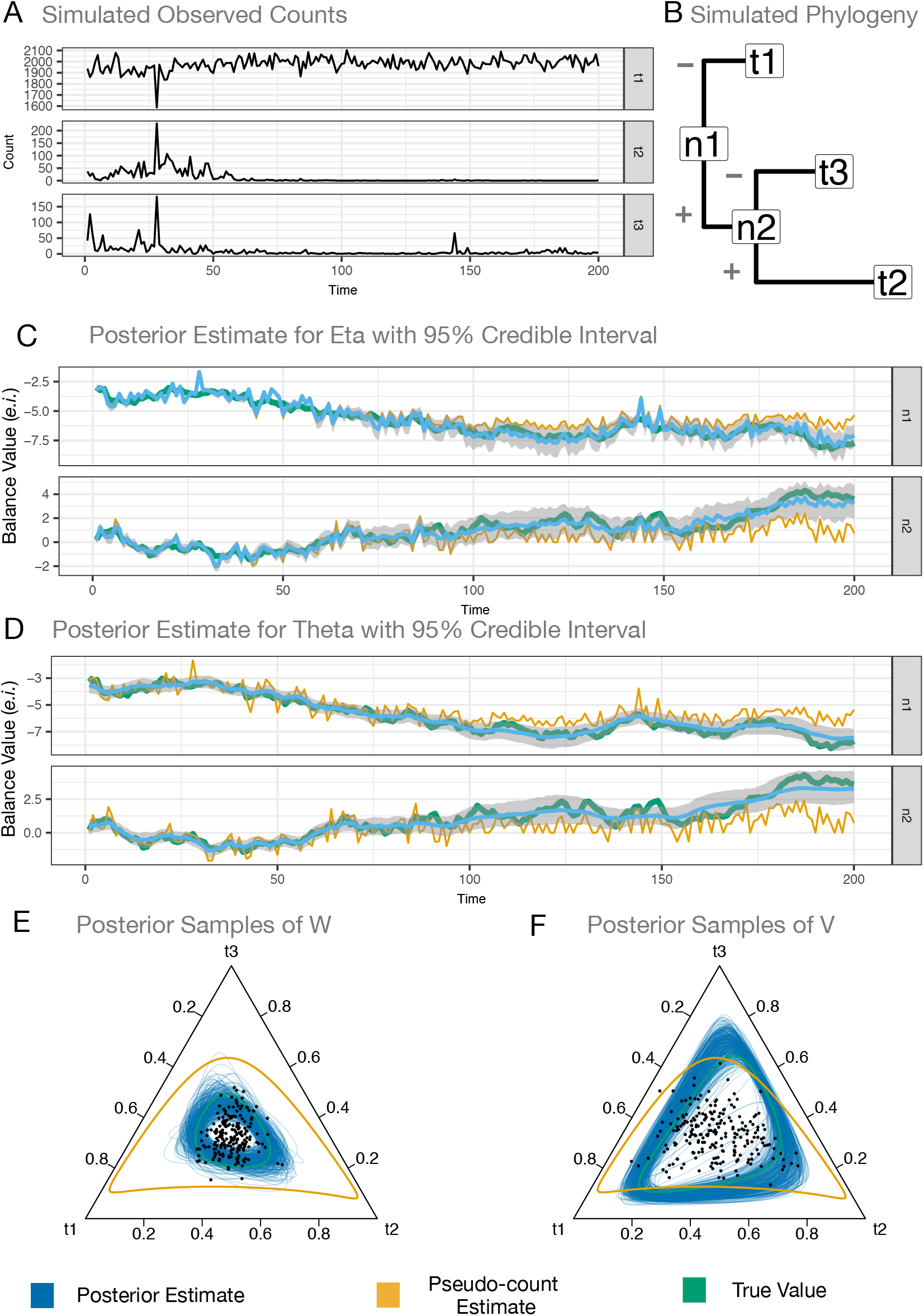
Analysis of a toy simulated microbial community demonstrates the advantages of accounting for technical noise and uncertainty due to counting. (**A**) A 3 taxon (t1, t2, t3) microbial community was simulated according to the likelihood model used to analyze the artificial gut dataset (*Methods*). Data from time-points 15, 16, and 20 were removed to simulate the effects of missing data on inferences. (**B**) A simulated phylogeny with annotated PhILR balances (n1, n2) used to analyze the simulated dataset. (+) and (−) refers to taxa in the numerator and denominator of associated balances. Pseudo-count based (PC) estimates for the multinomial parameters are obtained by adding 0.65 to all counts and then dividing each count by the sequencing depth of its associated sample and are shown as reference in (**C-D**). (**C**) Posterior mean and 95% credible interval for the multinomial parameters *η*. (**D**) Posterior mean and 95% credible interval for the unobserved microbial dynamics *θ*. PC estimates for the covariance of the multinomial parameters was obtained as the covariance of the first difference of the PC parameter estimates and are shown as reference in (**E-F**). 100 samples from the posterior distribution of the biological variation (*W*, **E**) and technical variation (*V*, **F**) depicted as 95% probability regions of the Logistic Normal distribution centered at the point in the simplex where each bacterial taxon is equally abundant. Black points represent the simulated biological variations (*w_t_*).

